# Simulation-based inference of evolutionary parameters from adaptation dynamics using neural networks

**DOI:** 10.1101/2021.09.30.462581

**Authors:** Grace Avecilla, Julie N. Chuong, Fangfei Li, Gavin Sherlock, David Gresham, Yoav Ram

## Abstract

The rate of adaptive evolution depends on the rate at which beneficial mutations are introduced into a population and the fitness effects of those mutations. The rate of beneficial mutations and their expected fitness effects is often difficult to empirically quantify. As these two parameters determine the pace of evolutionary change in a population, the dynamics of adaptive evolution may enable inference of their values. Copy number variants (CNVs) are a pervasive source of heritable variation that can facilitate rapid adaptive evolution. Previously, we developed a locus-specific fluorescent CNV reporter to quantify CNV dynamics in evolving populations maintained in nutrient-limiting conditions using chemostats. Here, we use the observed CNV adaptation dynamics to estimate the rate at which beneficial CNVs are introduced through *de novo* mutation and their fitness effects using simulation-based Bayesian likelihood-free inference approaches. We tested the suitability of two evolutionary models: a standard Wright-Fisher model and a chemostat growth model. We evaluated two likelihood-free inference algorithms: the well-established *Approximate Bayesian Computation with Sequential Monte Carlo* (ABC-SMC) algorithm, and the recently developed *Neural Posterior Estimation* (NPE) algorithm, which applies an artificial neural network to directly estimate the posterior distribution. By systematically evaluating the suitability of different inference methods and models we show that NPE has several advantages over ABC-SMC and that a Wright-Fisher evolutionary model suffices in most cases. Using our validated inference framework, we estimate the CNV formation rate at the *GAP1* locus in yeast as 10^−4.7^ -10^−4^ per cell division, and a selection coefficient of 0.04 - 0.1 per generation for *GAP1* CNVs in glutamine-limited chemostats. We experimentally validated our estimates using barcode lineage tracking and pairwise fitness assays. Our results are consistent with a high beneficial CNV supply rate that is 10-fold greater than the estimated rates of beneficial single-nucleotide mutations, explaining their outsized importance in rapid adaptive evolution. More generally, our study demonstrates the utility of novel simulation-based likelihood-free inference methods for inferring the rates and effects of evolutionary processes from empirical data.

## INTRODUCTION

Evolutionary dynamics are determined by the supply rate of beneficial mutations and their associated fitness effect. As the combination of these two parameters determines the overall rate of adaptive evolution, experimental methods are required for separately estimating them. The fitness effects of beneficial mutations are generally determined using competition assays and mutation rates are typically estimated using mutation accumulation or Luria-Delbrück fluctuation assays [1,2]. An alternative approach to estimating these evolutionary parameters entails quantifying the dynamics of adaptive evolution and using statistical inference methods to find parameter values that are consistent with the dynamics. Approaches to measure the dynamics of adaptive evolution, quantified as changes in the frequencies of beneficial alleles, have become increasingly accessible using either phenotypic makers or high-throughput DNA sequencing. Thus, inferential methods using adaptation dynamics data hold great promise for determining the underlying evolutionary parameters.

The combined fitness effects of all beneficial mutations define a distribution of fitness effects (DFE). Determining the DFE in a given condition is a central goal of evolutionary biology. Typically, beneficial mutations can occur at multiple loci and thus variance in the DFE reflects genetic heterogeneity. However, in some scenarios a single locus is the only gene in which beneficial mutations can occur, such as the case of mutations in the *β*-lactam gene conferring antibiotic resistance in bacteria [3]. In this case different mutations at the same locus confer differential beneficial effects resulting in a locus specific DFE. Thus, a DFE of beneficial mutations encompasses both allelic and locus heterogeneity.

Copy number variants (CNVs), defined as deletions or amplifications of genomic sequence, underlie rapid adaptive evolution in diverse scenarios ranging from niche adaptation to speciation [4–8]. CNVs are common in human populations [9–11], and pervasive among domesticated and wild populations of animals and plants [12–14], pathogenic and non-pathogenic microbes [15–18], and viruses [19–21]. CNVs can be both a driver and a consequence of cancers (reviewed in [22]). In the short term, CNVs may provide immediate fitness benefits by altering gene dosage. Over longer evolutionary timescales, CNVs can provide the raw material for the generation of evolutionary novelty through diversification of different gene copies [23].

Despite their prevalence, our understanding of the dynamics and reproducibility of CNVs in adaptive evolution is poor. Specifically, key evolutionary properties of CNVs, including their rate of formation and fitness effects, are largely unknown. As with other classes of genomic variation, CNVs are rare events occurring at sufficiently low frequencies to make empirical measurement difficult. Estimates of *de novo* CNV rates are derived from indirect and imprecise methods, and even when genome-wide mutation rates are directly quantified by mutation accumulation studies and whole-genome sequencing, estimates depend on both genotype and condition [2] and vary by orders of magnitude. Reported frequencies of duplications per locus per generation range from 10^−6^ to 10^−2^ in *Escherichia coli* and *Salmonella [24–29]*, 10^−6^ to 10^−4^in *Drosophila [30,31]*, 10^−7^ to 10^−5^ in human sperm [32,33], and 10^−12^ to 10^−6^ in the yeast *Saccharomyces cerevisiae* [34–38]. Reported rates of large scale duplications and aneuploidy in *S. cerevisiae* are 10^−5^ to 10^−4^ per cell per division [39–41].

There is also considerable evidence for heterogeneity in the CNV rate between loci, as factors including local sequence features, transcriptional activity, genetic background, and the external environment may impact the mutation spectrum. For example, there is evidence that CNVs occur at a higher rate near certain genomic features, such as repetitive elements [42], tRNA genes [43], origins of replication [44], and replication fork barriers [45]. Studies investigating the *CUP1* locus in budding yeast have shown that rates of amplification increase in response to transcriptional activation of *CUP1*, which is environmentally determined *[46,47]*.

Ploidy and diverse molecular mechanisms likely also impact CNV formation rates. Rates of aneuploidy, which result from nondisjunction errors, are higher in diploid yeast than haploid yeast, and chromosome gains are more frequent than chromosome losses [40]. Stress-induced mutagenesis, common in bacterial and cancer cells, may bias the mutational spectrum toward CNV because it activates mutagenic break repair, which can lead to CNV formation by microhomology-mediated break induced replication (reviewed in [48,49]). A recent study in yeast found that while overall mutation rates decrease with decreasing growth rate, CNV formation rates increase in slow-growing cells [50]. In human cells, hypoxia, but not other stressors, induces site-specific amplification that is also commonly observed in tumors [51].

The fitness effects of CNVs vary depending on gene content, genetic background and the environment. In evolution experiments in many systems, CNVs arise repeatedly in response to strong selection [52–60], suggesting they have strong beneficial fitness effects. Several of these studies measured fitness of clonal isolates containing CNVs, and reported selection coefficients of -0.11 to 0.6 [53,60,61]. However, the fitness of lineages containing CNVs varies between isolates even within single studies, which could be due to additional heritable variation or to differences in fitness between different types of CNVs (e.g. aneuploidy vs. single-gene amplification).

Due to the challenge of empirically measuring rates and effects of beneficial mutations across many genetic backgrounds, conditions, and types of mutations, researchers have attempted to infer these parameters from population-level data using evolutionary models and Bayesian inference [62–64]. This approach has several advantages. First, model-based inference provides estimations of interpretable parameters and the opportunity to compare multiple models. Second, the degree of uncertainty associated with a point estimate can be quantified. Third, a posterior distribution over model parameters allows exploration of parameter combinations that are consistent with the observed data, and posterior distributions can provide insight into certain relationships between parameters [65]. Fourth, posterior predictions can be generated using the model and either compared to the data or used to predict the outcome of differing scenarios.

Standard Bayesian inference requires a likelihood function, which gives the probability of obtaining the observed data given some values of the model parameters. However, for many evolutionary models, such as the Wright-Fisher model, the likelihood function is analytically or computationally intractable. Likelihood-free simulation-based Bayesian inference methods that bypass the likelihood function, such as *Approximate Bayesian Computation* (ABC; [66]), have been developed and used extensively in population genetics [67], ecology and epidemiology [68,69], and cosmology [70]. The simplest form of likelihood-free inference is rejection-ABC [71,72], in which model parameter proposals are sampled from a prior distribution, simulations are generated based on those parameter proposals, and simulated data are compared to empirical observations using a summary and distance function. Proposals that generate simulated data with a distance less than a defined tolerance threshold are considered samples from the posterior distribution and can therefore be used for its estimation. Efficient sampling methods have been introduced, namely MCMC [73] and SMC [74], that iteratively select proposals based on previous parameters samples so that regions of the parameter space with higher posterior density are explored more often. A shortcoming of ABC is that it requires summary and distance functions, which may be difficult to choose appropriately and compute efficiently, especially when using high-dimensional or multi-modal data, although methods have been developed to address this challenge [67,75,76]. Recently, new inferential methods have been introduced that directly approximate the likelihood or the posterior density function using *deep neural density estimators*—neural networks that approximate density functions. These methods, which have recently been used in neuroscience [65], population genetics [77], and cosmology [78], forego the summary and distance functions, can use data with higher dimensionality, and perform inference more efficiently [65,78,79]. However, neural network-based inference methods have not previously been applied to experimental evolution.

Previously, we developed a fluorescent CNV reporter system in the budding yeast, *Saccharomyces cerevisiae*, to quantify the dynamics of *de novo* CNVs during adaptive evolution [61]. Using this system we quantified CNV dynamics at the *GAP1* locus, which encodes a general amino acid permease, in nitrogen-limited chemostats for over 250 generations in multiple independent populations (**Figure 1A**). We found that *GAP1* CNVs reproducibly arise early and sweep through the population. By combining the *GAP1* CNV reporter with barcode lineage tracking and whole-genome sequencing we found that 10^2^–10^4^ independent CNV-containing lineages comprising diverse structures compete within populations.

**Figure 1.**
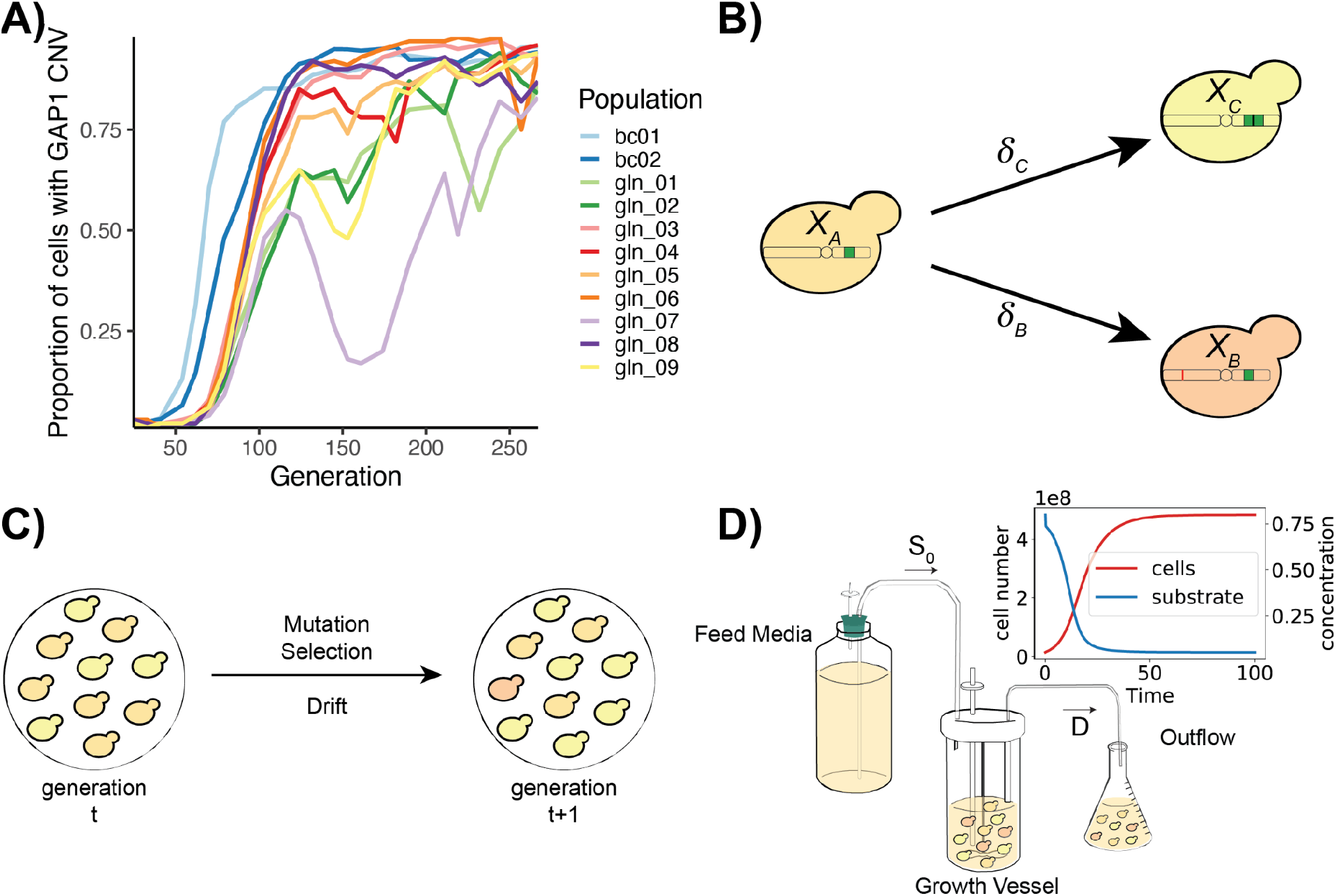
Empirical data and evolutionary models. **A)** Estimates of the proportion of cells with *GAP1* CNVs for eleven *S. cerevisiae* populations containing either a fluorescent *GAP1* CNV reporter (gln_01 - gln_09) or a fluorescent *GAP1* CNV reporter and lineage tracking barcodes (bc01 and bc02) evolving in glutamine-limited chemostats, from [61]. **B)** In our models, cells with the ancestral genotype (*X*_*A*_) can give rise to cells with a *GAP1* CNV (*X*_*C*_) or other beneficial mutation (*X*_*B*_) at rates *δ*_C_ and *δ*_B_, respectively. **C)** The Wright-Fisher model has discrete, non-overlapping generations and a constant population size. Allele frequencies in the next generation change from the previous generation due to mutation, selection, and drift. **D)** In the chemostat model, medium containing a defined concentration of a growth limiting nutrient (S_0_) is added to the culture at a constant rate. The culture, containing cells and medium, is removed by continuous dilution at rate *D*. Upon inoculation, the number of cells in the growth vessel increases and the limiting-nutrient concentration decreases until a steady state is reached (red and blue curves in inset). Within the growth vessel, cells grow in continuous, overlapping generations undergoing mutation, selection, and drift.

In this study, we estimate the formation rate and fitness effect of *GAP1* CNVs. We tested both ABC-SMC [80] and a neural density estimation method, NPE [81], using a classical Wright-Fisher model [82] and a chemostat model [83]. Using simulated data we tested the utility of the different evolutionary models and inference methods. We find that NPE has better performance than ABC-SMC. Although a more complex model has improved performance, the simpler and more efficient Wright-Fisher model is appropriate in most scenarios. We validated our approach by comparison to empirical estimates using lineage tracking and pairwise fitness assays. We estimate that beneficial *GAP1* CNVs are introduced at a rate of 10^−4.7^ -10^−4^ per cell division, and have a selection coefficient of 0.04 - 0.1 per generation. NPE is likely to be a useful method for inferring evolutionary parameters across a variety of scenarios, providing a powerful approach for combining experimental and computational methods.

## RESULTS

In a previous experimental evolution study, we quantified the dynamics of *de novo* CNVs in nine populations using a prototrophic yeast strain containing a fluorescent *GAP1* CNV reporter. [61]. Populations were maintained in glutamine-limited chemostats for over 250 generations and sampled every 8-20 generations (25 time points in total) to determine the proportion of cells containing a *GAP1* CNV using flow cytometry (populations gln_01-gln_09 **Figure 1A**). In the same study, we also performed two replicate evolution experiments using the fluorescent *GAP1* CNV reporter and lineage-tracking barcodes quantifying the proportion of the population with a *GAP1* CNV at 32 time points (populations bc01-bc02 in **Figure 1A**) [61]. We used interpolation to match timepoints between these two experiments (**Methods**) resulting in a dataset comprising the proportion of the population with a *GAP1* CNV at 25 timepoints in 11 replicate evolution experiments. In this study, we test whether the observed dynamics of CNV-mediated evolution provide a means of inferring the underlying evolutionary parameters.

### Overview of evolutionary models

We tested two models of evolution: the classical Wright-Fisher model [82] and a specialized chemostat model [83]. In both models, we start with an isogenic population in which *GAP1* CNV mutations occur at a rate *δ*_C_ and other beneficial mutations occur at rate *δ*_B_ (**Figure 1B**).

Previously, it has been shown that in populations in which clonal interference is common, adaptive dynamics are determined by broad properties of the distribution of fitness effects, and not its exact shape, and thus a single effective selection coefficient may be sufficient to model evolutionary dynamics [62]. Therefore, we focus on beneficial mutations and assume a single selection coefficient for each class of mutation. In our simulations, cells can acquire only a single beneficial mutation, either a CNV at *GAP1* or some other beneficial mutation (i.e. SNV, transposition, diploidization, or CNV at another locus). In all simulations (except for sensitivity analysis, see *Inference from empirical GAP1 dynamics*), the mutation rate of beneficial mutations other than *GAP1* CNVs was fixed at *δ*_B_=10^−5^ per genome per cell division and the selection coefficient was fixed at *s*_*B*_=0.001, based on estimates from previous experiments using yeast in several conditions [84–86]. Our goal was to infer the *GAP1* CNV mutation rate, *δ*_C_, and *GAP1* CNV selection coefficient, *s*_*C*_.

The two evolutionary models have several unique features. In the Wright-Fisher model, the population size is constant and each generation is discrete. Therefore, genetic drift is efficiently modeled using multinomial sampling (**Figure 1C**). In the chemostat model, fresh medium is added to the growth vessel at a constant rate and medium and cells are removed from the growth vessel at the same rate resulting in continuous dilution of the culture (**Figure 1D**). Individuals are randomly removed from the population through the dilution process, regardless of fitness, in a manner analogous to genetic drift. In the chemostat model, we start with a small initial population size and a high initial concentration of the growth-limiting nutrient. Following inoculation, the population size increases and the growth-limiting nutrient concentration decreases until a steady state is attained that persists throughout the experiment. As generations are continuous and overlapping in the chemostat model, we use the Gillespie algorithm with τ-leaping [87] to simulate the population dynamics. Growth parameters in the chemostat are based on experimental conditions during the evolution experiments [61] or taken from the literature (Table 1).

**Table 1.**
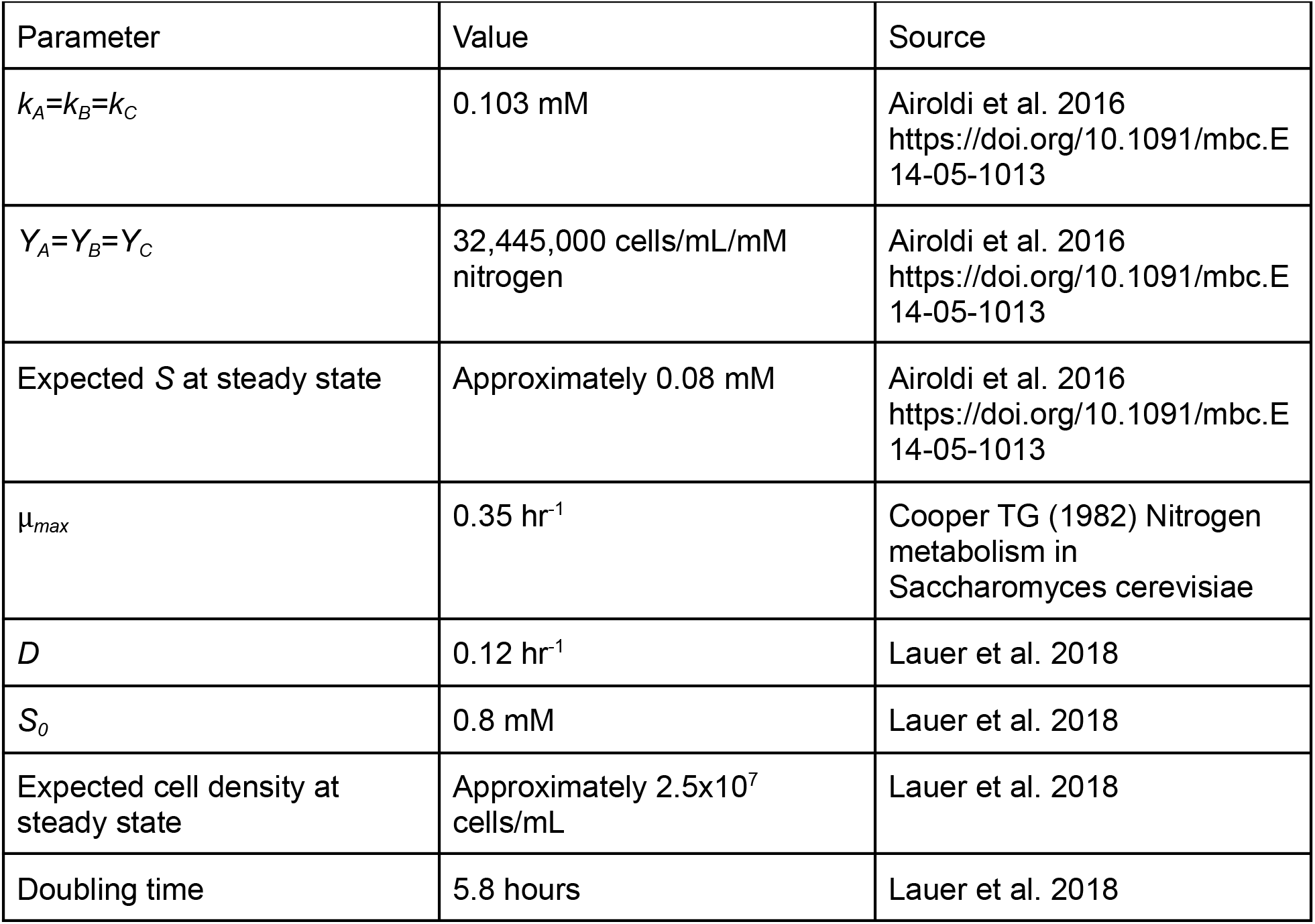
Chemostat parameters

### Overview of inference strategies

We tested two likelihood-free Bayesian methods for joint inference of the *GAP1* CNV mutation rate and the *GAP1* CNV fitness effect: Approximate Bayesian Computation with Sequential Monte Carlo (ABC-SMC) [74] and Neural Posterior Estimation (NPE) [88–90]. We used the proportion of the population with a *GAP1* CNV at 25 time points as the observed data (**Figure 1A)**. For both methods, we defined a log-uniform prior distribution for the CNV mutation rate ranging from 10^−12^ to 10^−3^ and a log-uniform prior distribution for the selection coefficient ranging from 10^−4^ to 0.4.

We applied ABC-SMC (**Figure 2A**), implemented in the Python package *pyABC* [80]. We used an adaptively weighted Euclidean distance function to compare simulated data to observed data. Thus, the distance function adapts over the course of the inference process based on the amount of variance at each time point [91]. The number of samples drawn from the proposal distribution during each iteration of SMC was adaptively determined based on the shape of the current posterior distribution [92] with a maximum number of samples per iteration set based on the total simulation budget (see **Methods**). We performed multiple iterations until either the acceptance threshold (ε = 0.002) was reached or until ten iterations had been completed.

**Figure 2.**
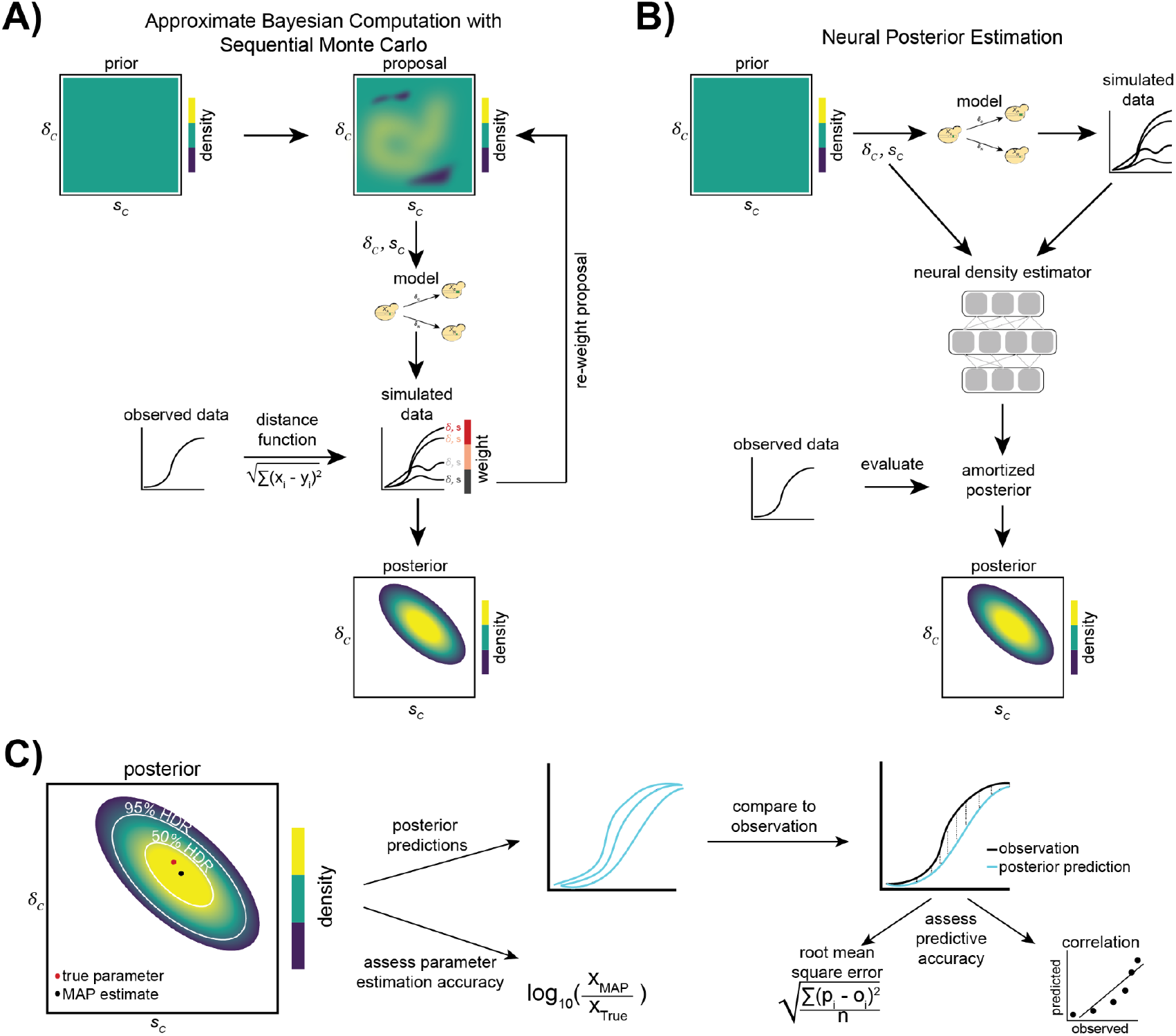
Inference strategies. **A)** When using Approximate Bayesian Computation with Sequential Monte Carlo (ABC-SMC), in the first iteration a proposal for the parameters *δ*_C_ (*GAP1* CNV mutation rate) and *s*_*C*_ (*GAP1* CNV selection coefficient) is sampled from the prior distribution. Simulated data are generated using either a Wright-Fisher or chemostat model and the current parameter proposal. The distance between the simulated data and the observed data is computed, and the proposed parameters are weighted by this distance. These weighted parameters are used to sample the proposed parameters in the next iteration. Over many iterations, the weighted parameter proposals provide an increasingly better approximation of the posterior distribution of *δ*_C_ and *s*_*C*_ (adapted from [79]). **B)** In Neural Posterior Estimation (NPE), simulated data are generated using parameters sampled from the prior distribution. From the simulated data and parameters, a density-estimating neural network learns the joint density of the model parameters and simulated data (the “amortized posterior”). The network then evaluates the conditional density of model parameters given the observed data, thus providing an approximation of the posterior distribution of *δ*_C_ and *s*_*C*_ (adapted from [79] and [65].) **C)** Assessment of inference performance. The 50% and 95% highest density regions (HDRs) are shown on the joint posterior distribution with the true parameters and the *maximum a posteriori* (MAP) parameter estimates. We compare the true parameters to the estimates by generating posterior predictions (sampling parameters from the joint posterior distribution and using them to simulate frequency trajectories), which we compare to the observations using the root mean square error (RMSE) and the correlation coefficient.

We applied NPE (**Figure 2B**), implemented in the Python package *sbi* [81], and tested two specialized normalizing flows as density estimators: a *masked autoregressive flow* (MAF) [93] and a *neural spline flow* (NSF) *[94]*. The normalizing flow is used as a density estimator to “learn” an amortized posterior distribution, which can then be evaluated for specific observations. Thus, amortization allows for evaluation of the posterior for each new observation without the need to re-train the neural network. To test the sensitivity of our inference results on the set of simulations used to learn the amortized posterior, we trained three independent amortized networks with different sets of simulations generated from the prior distribution and compared our resulting posterior distributions for each observation. We refer to inferences made with the same amortized network as having the same “training set.”

### NPE outperforms ABC-SMC

To test the performance of each inference method and evolutionary model, we generated 20 simulated synthetic observations for each model over four combinations of CNV formation rates and selection coefficients, resulting in 40 synthetic observations. We refer to the parameters that generated the synthetic observation as the “true” parameters. For each synthetic observation we performed inference using each method three times. Inference was performed using the same evolutionary model as that used to generate the observation. We found that with NPE, using NSF as the density estimator was superior to using MAF, and therefore we report results using NSF in the main text (results using MAF are in **Supplementary Figure 1**). Overall, we find that the accuracy of parameter inference depends on the model, method, and combination of parameters.

For each inference procedure we plotted the joint posterior distribution with the 50% and 95% highest density regions (HDR) [95] demarcated (**Figure 2C**, Supplementary XX). The true parameters are expected to lie within these HDRs at least 50% and 95% of the time respectively. We also computed the marginal 95% highest density intervals (HDI) [95] using the marginal posterior distributions for the *GAP1* CNV selection coefficient and *GAP1* CNV formation rate. We found that the true parameters were within the 50% HDR in half or more of the tests (averaged over three training sets) across a range of parameter values with the exception of ABC-SMC applied to the Wright-Fisher model when the *GAP1* CNV formation rate (*δ*_C_=10^−7^) and selection coefficient (*s*_*C*_=0.001) were both low (**Figure 3A**). The true parameters were within the 95% HDR in 100% of tests (**Supplementary Files**). The width of the HDI is informative about the degree of uncertainty associated with the parameter estimation. The HDIs for both fitness effect and mutation rate tend to be smaller when inferring with NPE compared to ABC-SMC, and this advantage of NPE is more pronounced when the CNV formation rate is high (*δ*_C_=10^−5^) (**Figure 3B-C**).

**Figure 3.**
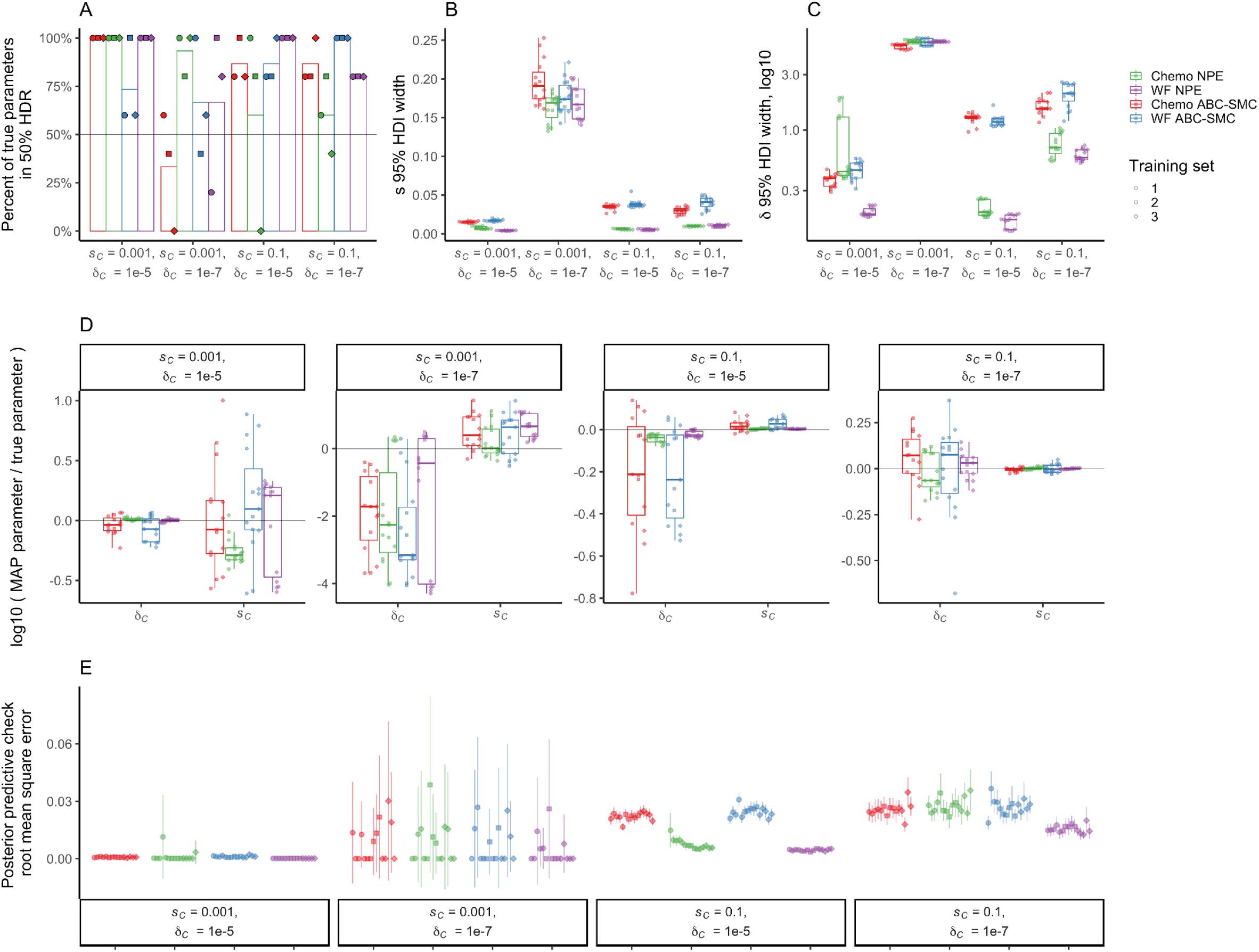
Performance assessment of inference methods using simulated synthetic observations. The figure shows the results of inference on five simulated synthetic observations using either the Wright-Fisher (WF) or chemostat (Chemo) model per combination of fitness effect *s*_*C*_ and mutation rate *δ*_C_. Simulations and inference were performed using the same model. For NPE, each training set corresponds to an independently amortized posterior distribution trained on a different set of 100,000 simulations, with which each synthetic observation was evaluated to produce a separate posterior distribution. For ABC-SMC, each training set corresponds to independent inference procedures on each observation using a total simulation budget of 10,000 per set and a stopping criteria of 10 iterations or ε <= 0.002, whichever occurs first. **A)** The percent of true parameters within the 50% HDR of the inferred posterior distribution. The bar height shows the average of three training sets. Horizontal line marks 50%. **B-C)** Distribution of widths of 95% highest density interval (HDI) of the posterior distribution of the fitness effect *s*_*C*_ (**B**) and CNV mutation rate *δ*_C_ (**C**), calculated as the difference between the 97.5 percentile and 2.5 percentile, for each separately inferred posterior distribution. **D)** Log-ratio of MAP estimate to true parameter for *s*_*C*_ and *δ*_C_. Note the different y-axis ranges. Grey horizontal line represents a log-ratio of zero, indicating an accurate MAP estimate. **E)** Mean and 95% confidence interval of RMSE of 50 posterior predictions compared to the synthetic observation from which the posterior was inferred.

We computed the *maximum a posteriori* (MAP) estimate of the *GAP1* CNV formation rate and selection coefficient by determining the mode (i.e. argmax) of the joint posterior distribution, and computed the log-ratio of the MAP relative to the true parameters. We find that the MAP estimate is close to the true parameter (i.e. the log-ratio is close to zero) when the selection coefficient is high (*s*_*C*_=0.1), regardless of the model or method, and much of the error is due to the mutation rate estimation error (**Figure 3D**). Generally, the MAP estimate is within an order of magnitude of the true parameter (i.e. the log-ratio is less than one), except when the formation rate and selection coefficient are both low (*δ*_C_=10^−7^, *s*_*C*_=0.001); in this case the formation rate was under-estimated up to four-fold and the selection coefficient was slightly over-estimated (**Figure 3D**). In some cases there are substantial differences in log-ratio due to training set using NPE; however, variation in log-ratio due to training set with NPE is usually less than the variation in the log-ratio when performing inference with ABC-SMC. Overall, the log-ratio tends to be closer to zero (i.e no difference) when using NPE (**Figure 3D**).

We performed posterior predictive checks by simulating *GAP1* CNV dynamics using the MAP estimates as well as 50 parameter values sampled from the posterior distribution (**Supplementary Files**). We computed both the root mean squared error (RMSE) and the correlation coefficient between posterior predictions and the observation to measure the prediction accuracy (**Figure 3E, Supplementary Figure 2**). We find that the RMSE posterior predictive accuracy of NPE is similar to, or better than, that of ABC-SMC (**Figure 3E**). The predictive accuracy quantified using correlation was close to 1 for all cases except when *GAP1* CNV formation rate and selection coefficient are both low (*s*_*C*_=0.001 and *δ*_C_=10^−7^) (**Supplementary Figure 2**).

We performed model comparison using both AIC (Akaike information criterion), computed using the MAP estimate, and WAIC (widely applicable information criterion), computed over the entire posterior distribution [96]. Lower values imply higher predictive accuracy and a difference of 2 is a standard threshold for model comparison (**Supplementary Figure 3**) [97]. We find similar results for both criteria: NPE with either model have similar values, though the value for Wright-Fisher is sometimes slightly lower than the value for the chemostat model. When *s*_*C*_=0.1, the value for NPE is consistently lower than for ABC-SMC, with a difference greater than 2. When *δ*_C_=10^−5^ and *s*_*C*_=0.001, the value for NPE with the Wright-Fisher model is lower than that for ABC-SMC with a difference of greater than 2, while the NPE with the chemostat model is not. The difference between any combination of model and method was less than 2 for *δ*_C_=10^−5^ and *s*_*C*_=0.001. Therefore, NPE is similar or better than ABC-SMC using either evolutionary model and for all tested combinations of *GAP1* CNV formation rate and selection coefficient.

We performed NPE using 10,000 or 100,000 simulations to train the neural network and found that increasing the number of training simulations did not substantially reduce the estimation error, but did tend to decrease the width of the 95% HDIs for both parameters (**Supplementary Figure 4**). Similarly, we performed ABC-SMC with the Wright-Fisher model with per observation budgets of 10,000 and 100,000 parameter samples (i.e. “particles” or “population size”), and found that increasing the budget decreased the estimation error and widths of the 95% HDIs for both parameters (**Supplementary Figure 4**).

### The Wright-Fisher model is suitable for inference using chemostat dynamics

While the chemostat model is a more precise description of our evolution experiments, both the model itself and its computational implementation have some drawbacks. First, the model is a stochastic continuous-time model implemented using the **τ**-leap method [87]. In this method, time is incremented in discrete steps and the number of stochastic events that occur within that time step is sampled based on the rate of events and system state at the previous time step. For accurate stochastic simulation, event rates and probabilities must be computed at each time step, and time steps must be sufficiently small. This incurs a heavy computational cost as time steps are considerably smaller than one generation, which is the time step used in the simpler Wright-Fisher model. Moreover, the chemostat model itself has additional parameters compared to the Wright-Fisher model, which must be experimentally measured or estimated.

The Wright-Fisher model is more general and more computationally efficient than the chemostat model. Therefore, we investigated if it can be used to perform accurate inference with NPE on synthetic observations generated by the chemostat model. By assessing how often the true parameters fell within the HDRs, we found that the Wright-Fisher is a good-enough approximation of the full chemostat dynamics when selection is weak (*s*_*C*_ = 0.001) (**Supplementary Figure 5**), and it performs similarly to the chemostat model in parameter estimation accuracy (**Figure 4A-B**). The Wright-Fisher is less suitable when selection is strong (*s*_*C*_ = 0.1), as the true parameters do not fall within the 50% or 95% HDR (**Supplementary Figure 5**). Nevertheless, estimation of the selection coefficient remains accurate, and the difference in estimation of the formation rate is less than an order of magnitude, with a 3-5-fold overestimation (MAP log-ratio between 0.5 and 0.7) (**Figure 4C-D**).

**Figure 4.**
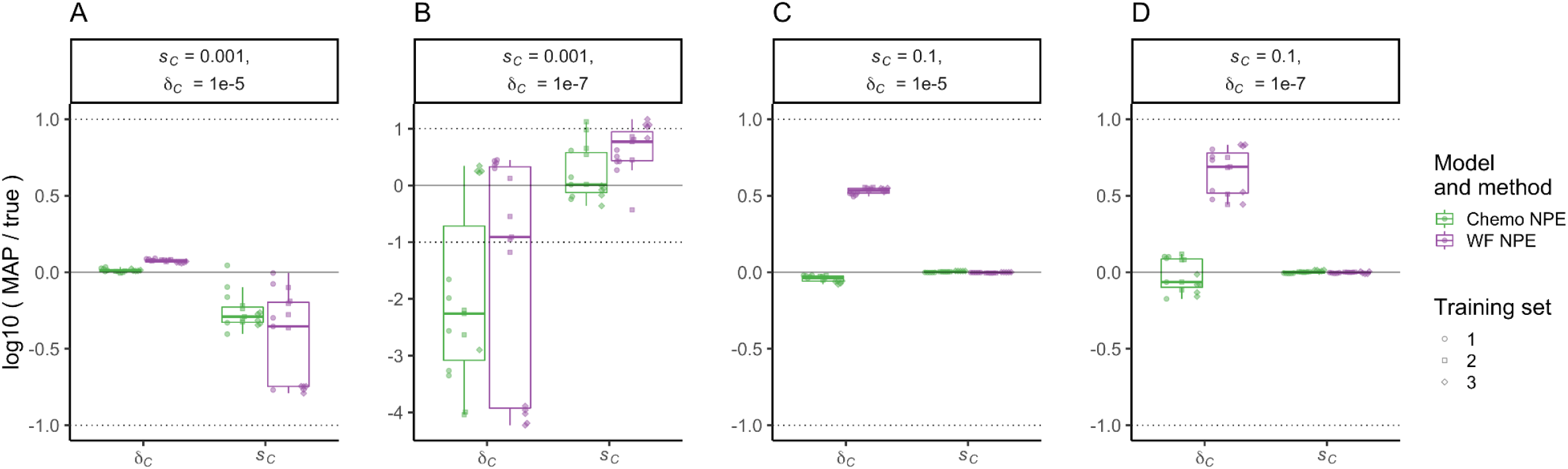
Inference with Wright-Fisher model from chemostat dynamics. The figure shows results of inference using NPE and either the Wright-Fisher (WF) or chemostat (Chemo) model on five simulated synthetic observations generated using the chemostat model for different combinations of fitness effect *s*_*C*_ and formation rate *δ*_C_. Boxplots and markers show the log-ratio of MAP estimate to true parameters for *s*_*C*_ and *δ*_C_. Horizontal solid line represents a log-ratio of zero, indicating an accurate MAP estimate; dotted lines indicate an order of magnitude difference between the MAP estimate and the true parameter.

### Inference using a set of observations

Our empirical dataset includes eleven biological replicates of the same evolution experiment. Differences in the dynamics between independent replicates may be explained by an underlying distribution of fitness effects (DFE) rather than a single constant selection coefficient. It is possible to infer the DFE using all experiments simultaneously. However, inference of distributions from multiple experiments presents several challenges, common to other mixed-effects or hierarchical models [98]. Alternatively, individual values inferred from individual experiments could provide an approximation of the underlying DFE.

To test these two alternative strategies for inferring the DFE, we performed simulations in which we allowed for variation in the selection coefficient of *GAP1* CNVs for each population in a set of observations. We sampled eleven selection coefficients from a Gamma distribution with shape and scale parameters *α* and *β*, respectively, and an expected value *E*(*s*) = *αβ* [99], and then simulated a single observation for each sampled selection coefficient. As the Wright-Fisher model is a suitable approximation of the chemostat model (**Figure 4**), we used the Wright-Fisher model both for generating our observation sets and for parameter inference.

For the observation sets, we used NPE to either infer a single selection coefficient for each observation or to directly infer the Gamma distribution parameters *α* and *β* from all eleven observations. When inferring eleven selection coefficients, one for each observation in the observation set, we fit a Gamma distribution to eight of the eleven inferred values (**Figure 5**, green lines). When directly inferring the DFE, we used a uniform prior for *α* from 0.5 to 15 and a log-uniform prior for *β* from 10^−3^ to 0.8. We held out three experiments from the set of eleven and used a three-layer neural network to reduce the remaining eight observations to a five-feature summary statistic vector, which we then used as an embedding net [81] with NPE to infer the joint posterior distribution of *α, β*, and *δ*_C_ (**Figure 5**, blue lines). For each observation set, we performed each inference method three times, using different sets of eight experiments to infer the underlying DFE.

**Figure 5.**
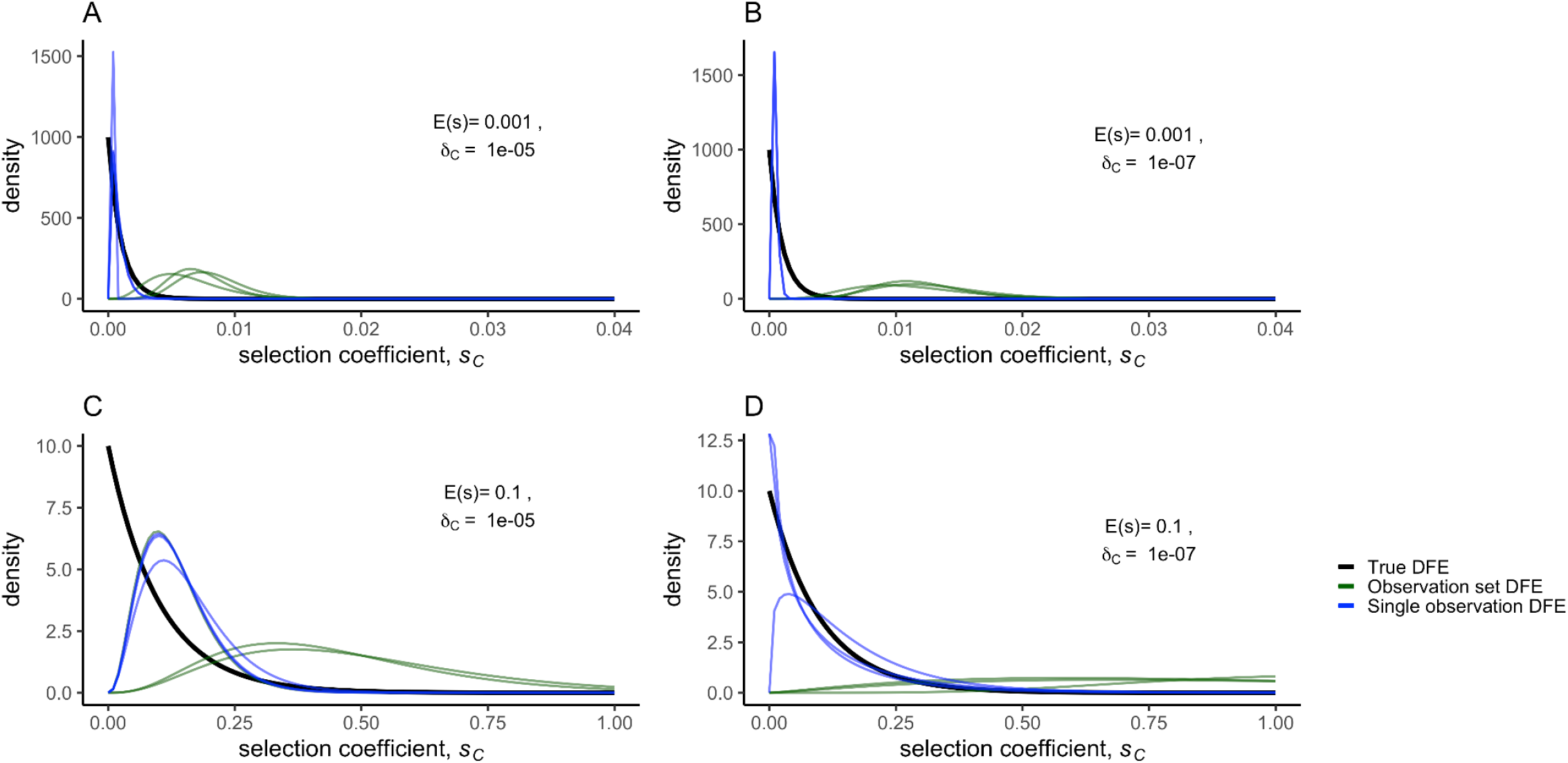
Inference of the distribution of fitness effects. A set of eleven simulated synthetic observations was generated from a Wright-Fisher model with CNV selection coefficients sampled from an exponential (Gamma with *α* = 1) distribution of fitness effects (true DFE; black curve). The MAP DFEs (observation set DFE, green curves) were directly inferred using three different subsets of eight out of eleven synthetic observations. We also inferred the selection coefficient for each individual observation in the set of eleven separately, and fit a Gamma distribution (single observation DFE, blue curves) to sets of eight inferred selection coefficients. All inferences were performed with NPE using the same amortized network to infer a posterior for each set of eight synthetic observations or each single observation. **A**) weak selection, high formation rate, **B)** weak selection, low formation rate, **C**) strong selection, high formation rate, **D**) strong selection, low formation rate.

We used Kullback–Leibler divergence to measure the difference between the true DFE and inferred DFE, and find that the inferred selection coefficients from the single experiments capture the underlying DFE as well or better than direct inference of the DFE from a set of observations for both *α* = 1 (an exponential distribution) and *α* = 10 (sum of ten exponentials) (**Figure 5)**. The only exception we found is when *α* = 10, *E*(*s*) = 0. 001, and *δ*_C_=10^−5^ (**Supplementary Figure 6, Supplementary Table 1**). We assessed the performance of inference from a set of observations using out-of-sample posterior predictive accuracy [96] and found that inferring *α* and *β* from a set of observations results in lower posterior predictive accuracy compared to inferring *s*_*C*_ from a single observation (**Supplementary Figure 7**). Therefore, estimating the DFE through inference of individual selection coefficients from each observation is superior to inference of the distribution from multiple observations.

### Inference from empirical evolutionary dynamics

To apply our approach to empirical data we inferred *GAP1* CNV selection coefficients and formation rates using eleven replicated evolutionary experiments in glutamine-limited chemostats [61] (**Figure 1A**) using NPE with both models. We performed posterior predictive checks, drawing parameter values from the posterior distribution, and found that *GAP1* CNV were predicted to increase in frequency earlier and more gradually than is observed in our experimental populations (**Supplementary Figure 8**). This discrepancy is especially apparent in experimental populations that appear to experience clonal interference with other beneficial lineages (i.e. gln07, gln09). Therefore, we excluded data after generation 116, by which point CNVs have reached high frequency in the populations but do not yet exhibit the non-monotonic and variable behavior observed in later time points, and re-did the inference. The resulting posterior predictions are more similar to the observations in initial generations (average MAP RMSE for the eleven observations up to generation 116 is 0.06 when inference excludes late time points versus 0.13 when inference includes all time points). Furthermore, the overall RMSE (for observations up to generation 267) was not significantly different (average MAP RMSE is 0.129 and 0.126 when excluding or including late time points, respectively; **Supplementary Figure 9**). Restricting the analysis to early time points did not dramatically affect estimates of *GAP1* CNV selection coefficient and formation rate, but it did result in less variability in estimates between populations (i.e. independent observations) and some reordering of populations’ selection coefficients and formation rate relative to each other (**Supplementary Figure 10**). Thus, we focused on inference using data prior to generation 116.

The inferred *GAP1* CNV selection coefficients were similar regardless of model, with the range of MAP estimates for all populations between 0.04 and 0.1, whereas the range of inferred *GAP1* CNV formation rates was somewhat higher when using the Wright-Fisher model, 10^−4.1^-10^−3.4^, compared to the chemostat model, 10^−4.7^ -10^−4^ (**Figure 6A-B**). While there is variation in inferred parameters due to the training set, variation between observations is higher than variations between training sets (**Figure 6A-C**). Posterior predictions using the chemostat model, a fuller depiction of the evolution experiments, tend to have slightly lower RMSE than predictions using the Wright-Fisher model (**Figure 6C**). However, predictions using both models greatly resemble the true *GAP1* CNV dynamics, especially in early generations (**Figure 6D**).

**Figure 6.**
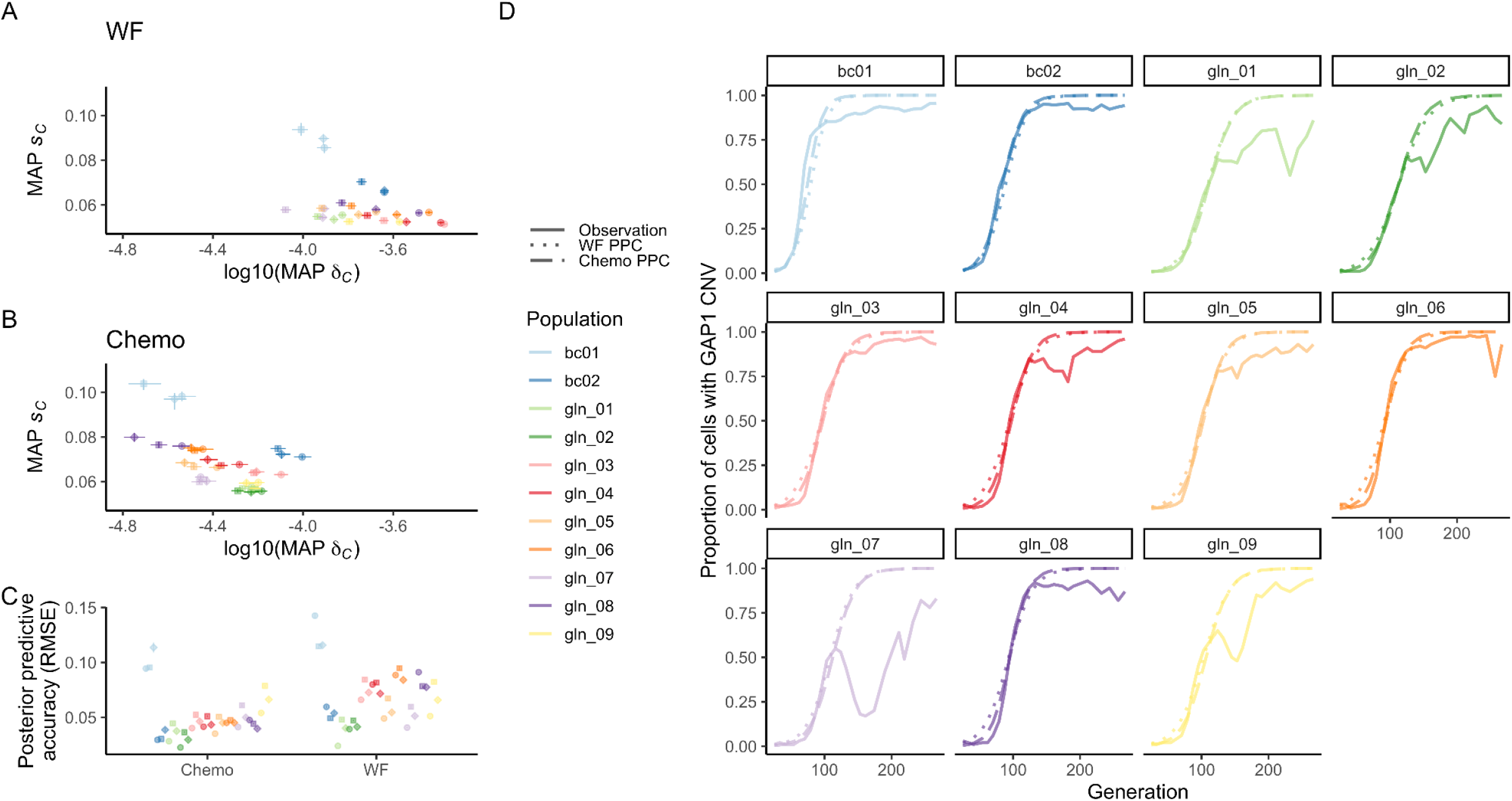
Inference of CNV formation rate and fitness effect from empirical evolutionary dynamics. The inferred MAP estimate and 95% highest density intervals (HDI) for fitness effect *s*_*C*_ and formation rate *δ*_C_, using the **(A)** Wright-Fisher (WF) or **(B)** chemostat (Chemo) model and NPE for each experimental population from [61]. Inference performed with data up to generation 116, and each training set corresponds to three independent amortized posterior distributions estimated with 100,000 simulations. **C)** Mean and 95% confidence interval for RMSE of 50 posterior predictions compared to empirical observations up to generation 116. **D)** Proportion of the population with a *GAP1* CNV in the experimental observations (solid lines) and in posterior predictions using the MAP estimate from training set 1 shown in panels A and B with either the Wright-Fisher (dotted line) or chemostat (dashed line) model. Mutation rate and fitness effect of other beneficial mutations set to 10^−5^ and 10^−3^, respectively.

To test the sensitivity of these estimates, we also inferred the *GAP1* CNV selection coefficient and formation rate using the Wright-Fisher model in the absence of other beneficial mutations (*δ*_B_=0), when other beneficial mutations have an intermediate selection coefficient (*s*_*B*_=0.01, *δ*_B_=10^−5^), when other beneficial mutations have a high selection coefficient (*s*_*B*_=0.1, *δ*_B_=10^−5^), and when other beneficial mutations have a high selection coefficient but an even lower formation rate (*s*_*B*_=0.1, *δ*_B_=10^−7^). In general, perturbations to the rate and selection coefficient of other beneficial mutations did not alter the inferred *GAP1* CNV selection coefficient or formation rate. We found a single exception: when both the formation rate and fitness effect of other beneficial mutations is high (*s*_*B*_=0.1 and *δ*_B_=10^−5^), the *GAP1* CNV selection coefficient was approximately 1.6-fold higher and the formation rate was approximately 2-fold lower (**Supplementary Figure 11**); however, posterior predictions were poor for this set of parameter values (**Supplementary Figure 12**).

### Experimental confirmation of fitness effects inferred from adaptive dynamics

To validate the inferred selection coefficients, we used lineage tracking to estimate the distribution of fitness effects [100–102]. We used barseq on the entire evolving population at multiple time points and labeled lineages that did and did not contain *GAP1* CNVs (**Figure 7A**). Using barcode trajectories to estimate fitness effects ([100]; see **Methods**), we identified 1,569 out of 80,751 lineages (1.94%) as adaptive in the bc01 population. A total of 1,513 (96.4%) adaptive lineages have a *GAP1* CNV (**Figure 7A**). Selection coefficients can be directly measured using competition assays by fitting a linear model to the log-ratio of the *GAP1* CNV strain and ancestral strain frequencies over time (**Figure 7B**). We isolated *GAP1* CNV containing clones from populations bc01 and bc02, determined their fitness (**Methods)**, and combined these estimates with previously reported selection coefficients for *GAP1* CNV containing clones isolated from populations gln01-gln09 [61] to determine the DFE.

**Figure 7.**
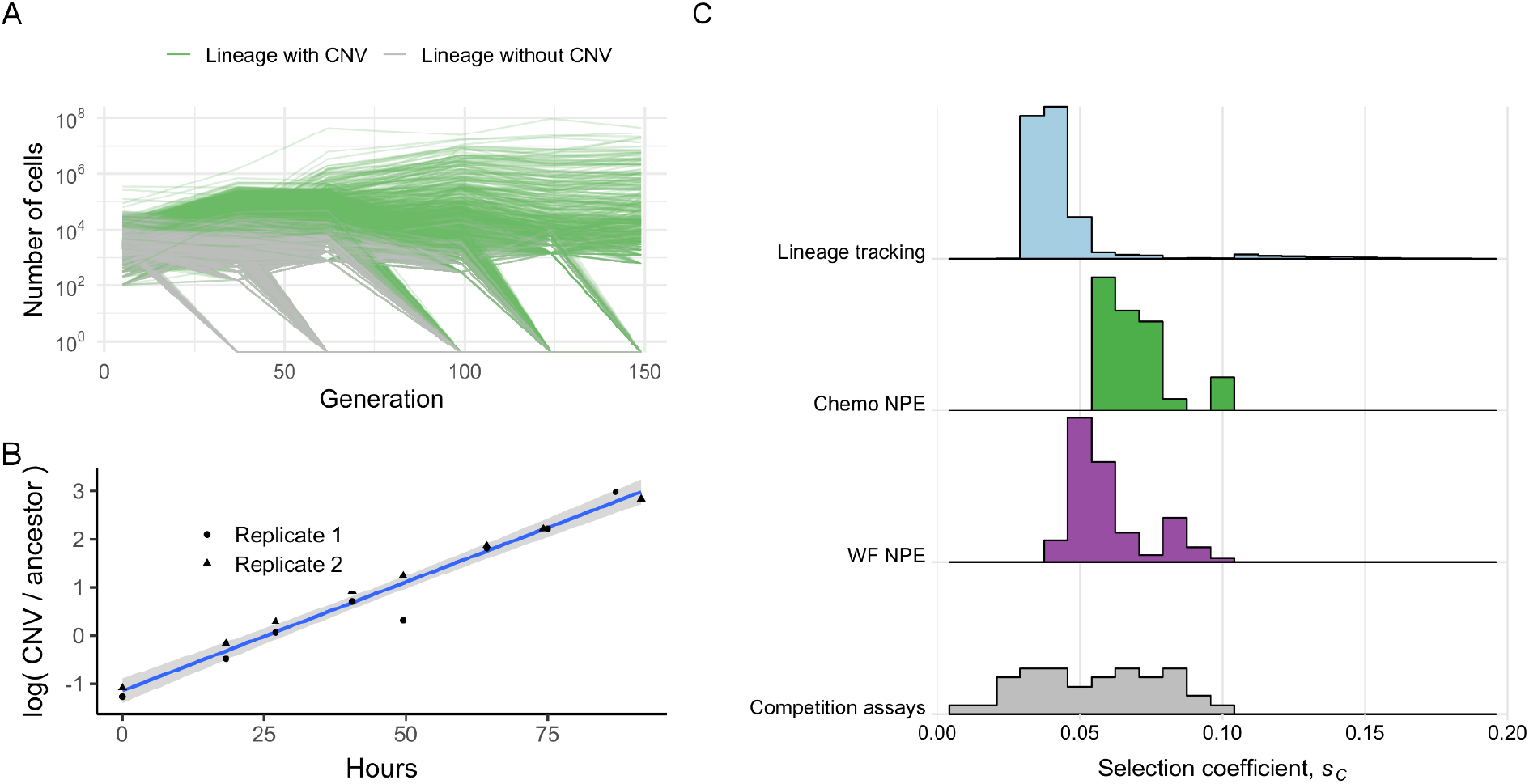
Comparison of DFE inferred using NPE, lineage-tracking barcodes, and competition assays. **A)** Barcode-based lineage frequency trajectories in experimental population bc01. Lineages with (green) and without (grey) *GAP1* CNVs are shown. **B)** Two replicates of a pairwise competition assay for a single *GAP1* CNV containing clone isolated from an evolving population. The selection coefficient for the clone is estimated from the slope of the linear model (blue line) and 95% CI (gray). **C)** The distribution of fitness effects for all beneficial *GAP1* CNVs inferred from eleven populations using NPE and the Wright-Fisher (WF; purple) and chemostat (Chemo; green) models compared with the DFE inferred from barcode frequency trajectories in the bc01 population (light blue) and the DFE inferred using pairwise competition assays with different *GAP1* CNV containing clones (grey).

The DFE for adaptive *GAP1* CNV lineages in bc01 inferred using lineage-tracking barcodes and the DFE from pairwise competition assays share similar properties to the distribution inferred using NPE from all experimental populations (**Figure 7C, Supplementary Figure 13**). Thus, our inference framework using CNV adaptation dynamics is a reliable estimate of the DFE estimated using laborious and more costly experimental methods.

## DISCUSSION

In this study we tested the application of simulation-based inference to the determination of key evolutionary parameters from observed adaptive dynamics in evolution experiments. We focused on the role of CNVs in adaptive evolution using experimental data in which we quantified the population frequency of *de novo* CNVs at a single locus using a fluorescent CNV reporter. The goal of our study was to determine the appropriate computational framework for simulation-based inference and to apply it to estimate the *GAP1* CNV selection coefficient and formation rates in experimental evolution using nitrogen-limited chemostats.

Our study yielded several important methodological findings. Using synthetic data we tested two different model-based algorithms for joint inference of evolutionary parameters, the effect of different evolutionary models on inference performance, and how best to determine a distribution of fitness effects using multiple experiments. We find that the neural-network-based algorithm NPE outperforms ABC-SMC regardless of evolutionary model. Although a more complex evolutionary model better describes the evolution experiments performed in chemostats, we find that a standard Wright-Fisher model is a sufficient approximation for inference using NPE. Finally, although it is possible to perform joint inference on multiple independent experimental observations to infer a distribution of fitness effects, we find that inference performed on individual experiments and post facto estimation of the distribution more accurately captures the underlying distribution of fitness effects.

Despite being originally developed to analyze population-genetic data [71–74], likelihood-free methods are uncommon in the field of experimental evolution. Previous studies that applied likelihood-free inference to results of evolutionary experiments differ from our study in various ways [62–64]. First, they used serial-dilution rather than chemostat experiments. Second, most focused on all beneficial mutations, whereas we categorize beneficial mutations into two categories: *GAP1* CNVs and all other beneficial mutations; thus, they used an evolutionary model with a single process generating genetic variation whereas our study includes two such processes, but focuses inference on our mutation type of interest. Third, we used two different evolutionary models: the Wright-Fisher model, a standard model in evolutionary genetics, and a chemostat model. The latter is more realistic but also more computationally demanding. Fourth, previous studies applied relatively simple rejection-ABC methods [62–64,99]. We applied two modern approaches: ABC with sequential Monte Carlo sampling [74], which is a computationally efficient algorithm for Bayesian inference, using an adaptive distance function [91]; and NPE [88–90] with *NSF* [94]. NPE approximates an amortized posterior distribution from simulations. Thus, it can be more efficient than ABC-SMC, as it can estimate a posterior distribution for new observations without requiring additional training. Importantly, our study is the first, to our knowledge, to use neural density estimation to apply likelihood-free inference to experimental evolution data.

Our application of simulation-based inference yielded new insights into the role of CNVs in adaptive evolution. We estimated *GAP1* CNV formation rate and selection coefficient from empirical population-level adaptive evolution dynamics and found that *GAP1* CNVs form at a rate of 10^−3.5^-10^−4.5^ per generation (approximately 1 in 10,000 cell divisions) and have selection coefficients of 0.05-0.1 per generation. We experimentally validated our inferred fitness estimates using barcode lineage tracking and pairwise competition assays and showed that simulation-based inference is in good agreement with the two different experimental methods. The formation rate that we have determined for *GAP1* CNVs is remarkably high. Locus-specific CNV formation rates are extremely difficult to determine and fluctuation assays have yielded estimates from 10^−12^ to 10^−6^ ^[34–38]^. Mutation accumulation studies have yielded genome-wide CNV rates of about 10^−5^ ^[39–41]^. Our estimated rate of *GAP1* CNV formation is higher than we would expect based on these previous estimates. We posit two possible explanations for this high rate: 1) CNVs at the *GAP1* locus may be deleterious in most conditions, including the putative non-selective conditions used for mutation-selection experiments, and therefore underestimated due to negative selection; and 2) under nitrogen-limiting selective conditions, in which *GAP1* expression levels are extremely high, a mechanism of induced CNV formation may operate that increases the rate at which they are generated, as has been shown at other loci in the yeast genome [46,47] Empirical validation of the inferred rate of *GAP1* CNV formation awaits experimental confirmation.

This simulation-based inference approach can be readily extended to other evolution experiments. In this study we performed inference of parameters for a single type of mutation. This approach could be extended to infer the rates and effects of multiple types of mutations simultaneously. For example, instead of assuming a rate and selection coefficient for other beneficial mutations and performing ex post facto analyses looking at the sensitivity of inference of *GAP1* CNV parameters in other beneficial mutation regimes, one could simultaneously infer parameters for both of these types of mutations. As shown using our barcode-sequencing data, many CNVs arise during adaptive evolution, and previous studies have shown that CNVs have different structures and mechanisms of formation [61,103]. Inferring a single effective selection coefficient and formation rate is a current limitation of our study that could be overcome by inferring rates and effects for different classes of CNVs (e.g, aneuploidy vs tandem duplication). Inspecting conditional correlations in posterior distributions involving multiple types of mutations has the potential to provide insights into how interactions between different classes of mutations shape evolutionary dynamics.

The approach could also be applied to CNV dynamics at other loci, in different genetic backgrounds, or in different media conditions. Furthermore, this approach could be used to infer formation rates and selection coefficients for other types of mutations; the empirical data required is simply the proportion of the population with a given mutation type over time, which can efficiently be determined using a phenotypic marker, or similar quantitative data such as whole-genome whole-population sequencing. Evolutionary models could be extended to more complex evolutionary scenarios including changing population sizes, fluctuating selection, and changing ploidy and reproductive strategy, with an ultimate goal of inferring their impact on a variety of evolutionary parameters and predicting evolutionary dynamics in complex environments and populations.

## METHODS

All source code for performing the analyses and reproducing the figures is available at https://github.com/graceave/cnv_sims_inference. All of the data can be found at https://osf.io/e9d5x/.

### Evolutionary models

We modeled the adaptive evolution from an isogenic asexual population with frequencies X_A_ of the ancestral (or wild-type) genotype, X_C_ of cells with a *GAP1* CNV, and X_B_ of cells with a different type of beneficial mutation. Ancestral cells can gain a *GAP1* CNV or another beneficial mutation at rates *δ*_C_ and *δ*_B_, respectively. Therefore, the frequencies of cells of different genotypes after mutation are

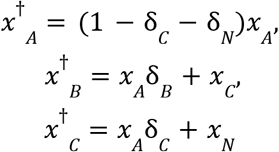

For simplicity, this model neglects cells with multiple mutations, which is reasonable for short timescales, such as those considered here.

In the discrete time Wright-Fisher model, the change in frequency due to natural selection is modeled by

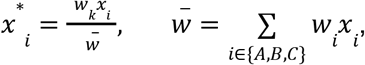

where w_i_ is the relative fitness of cells with genotype i, and 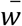 is the population mean fitness relative to the ancestral type. Relative fitness is related to the selection coefficient by

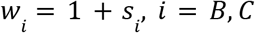

The change in due random genetic drift is given by

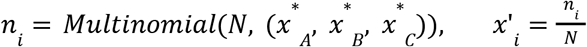

where N is the population size. In our simulations N=3.3×10^8^, the effective population size in the chemostat populations in our experiment (see **Determining effective population size in the chemostat**).

The chemostat model starts with a population size 1.5×10^−7^ and the concentration of the limiting nutrient in the growth vessel, *S*, is equal to the concentration of that nutrient in the fresh media, *S*_*0*_. During continuous culture, the chemostat is continuously diluted as fresh media flows in and culture media and cells are removed at rate *D*. During the initial phase of growth, the population size grows and the limiting nutrient concentration is reduced until a steady state is attained at which the population size and limiting nutrient concentration are maintained indefinitely. We extended the model for competition between two haploid clonal populations for a single growth-limiting resource in a chemostat from [83] to three populations such that

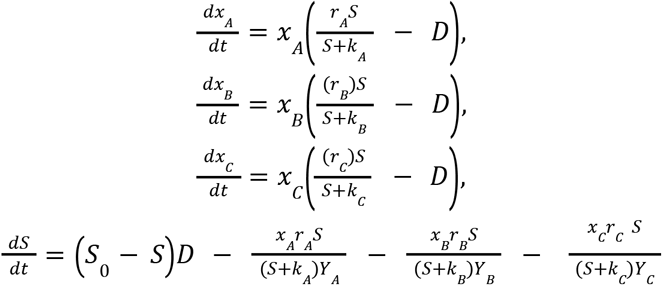

*Y*_*i*_ is the culture yield of strain i per mole of limiting nutrient. *r*_*A*_ is the Malthusian parameter, or intrinsic rate of increase, for the ancestral strain, and in the chemostat literature is frequently referred to as μ_*max*_, the maximal growth rate. The growth rate in the chemostat, μ, depends on the the concentration of the limiting nutrient with saturating kinetics 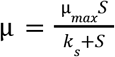. *k*_*i*_ is the substrate concentration at half-maximal μ. *r*_*C*_ and *r*_*B*_ are the Malthusian parameters for strains with a CNV and strains with an other beneficial mutation, respectively, and are related to the ancestral Malthusian parameter and selection coefficient by [104]

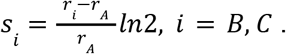

The values for the parameters used in the chemostat model are in Table 1.

We simulated continuous time in the chemostat using the Gillespie algorithm with τ-leaping. Briefly, we calculate the rates of ancestral growth, ancestral dilution, CNV growth, CNV dilution, other mutant growth, other mutant dilution, mutation from ancestral to CNV, and mutation from ancestral to other mutant. For the next time interval τ we calculated the number of times each event occurs during the interval using the Poisson distribution. The limiting substrate concentration is then adjusted accordingly. These steps repeat until the desired number of generations is reached.

For the chemostat model, we began counting generations after 48 hours, which is approximately the amount of time required for the chemostat to reach steady state, and when we began recording generations in [61].

### Determining the effective population size in the chemostat

In order to determine the effective population size in the chemostat, and thus the population size to use in with the Wright-Fisher model, we determined the conditional variance of the allele frequency in the next generation *p’* given the frequency in the current generation *p* in the chemostat. To do this, we simulated a chemostat population with two neutral alleles with frequencies *p* and *q* (*p+q=1*), which begin at equal frequencies, *p=q*. We allowed the simulation to run for 1,000 generations, recording the frequency *p* at every generation, excluding the first 100 generations to ensure the population is at steady state. We then computed the conditional variance *Var(p’*|*p)* in each generation and estimated the effective population size as (where *t=900* is the total number of generations)

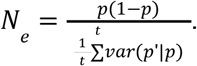

The estimated effective population size in our chemostat conditions is 3.3×10^8^, which is approximately two thirds of the census population size *N* when the chemostat is at steady state.

### Inference methods

For inference using single observations, we used the proportion of the population with a *GAP1* CNV at 25 time points as our summary statistics and defined a log-uniform prior for the mutation rate ranging from 10^−12^ to 10^−3^ and a log-uniform prior for the selection coefficient from 10^−4^ to 0.4.

For inference using sets of observation, we used a uniform prior for *α* from 0.5 to 15, a log-uniform prior for *β* from 10^−3^ to 0.8, and a log-uniform prior for the mutation rate ranging from 10^−12^ to 10^−3^. For use with NPE, we used a three layer sequential neural network with linear transformations in each layer and Rectified Linear Unit as the activation functions to encode the observation set into five summary statistics, which we then used as an embedding net with NPE.

We applied ABC-SMC implemented in the Python package *pyABC* [80]. For inference using single observations we used an adaptively weighted Euclidean distance function with the root mean square deviation as the scale function. For inference using a set of observations, we used the squared Euclidean distance as our distance metric. We used 100 samples from the prior for initial calibration before the first round, and simulation budgets of either 10,000 or 100,000 for both single observations and observation sets (i.e.10,000 single observations or 10,000 sets of 11 observations). For the 10,000 simulation budget, we started inference with 100 samples, had a maximum of 1,000 samples per round, and a maximum of ten rounds. For the 100,000 simulation budget, we started inference with 1,000 samples, had a maximum of 10,000 samples per round, and a maximum of ten rounds. The exact number of samples from the proposal distribution during each round of sampling were adaptively determined based on the shape of the current posterior distribution [92]. For inference of the posterior for each observation, we performed multiple rounds of sampling until either we reached the acceptance threshold ε <= 0.002 or ten rounds were performed.

We applied NPE implemented in the Python package *sbi* [81] using a *Masked Autoregressive Flow* (*MAF*) [93] or a a *neural spline flow (NSF) [94]* as a conditional density estimator that learns an amortized posterior density for single observations. We used either 10,000 or 100,000 simulations to train the network. To test the dependence of our results on the set of simulations used to learn the posterior, we trained three independent amortized networks with different sets of simulations generated from the prior and compared our resulting posterior distributions for each observation.

### Assessment of performance of each method with each model

To test each method, we simulated five populations for each combination of the following CNV mutation rates and fitness effects: *s*_*C*_=0.001 and *δ*_C_=10^−5^; *s*_*C*_=0.1 and *δ*_C_=10^−5^; *s*_*C*_=0.001 and *δ*_C_=10^−7^; *s*_*C*_=0.1 and *δ*_C_=10^−7^, for both the Wright-Fisher model and the chemostat model, resulting in 40 total simulated observations. We independently inferred the CNV fitness effect and mutation rate for each simulated observation three times.

We calculated the MAP estimate by first estimating a Gaussian kernel density estimate (KDE) using *SciPy* (*scipy*.*stats*.*gaussian_kde)* [105] with at least 1,000 parameter combinations and their weights drawn from the posterior distribution. We then found the maximum of the KDE (using *scipy*.*optimize*.*minimize* with the Nelder-Mead solver). We calculated the 95% highest density intervals for the MAP estimate of each parameter using *pyABC* (*pyabc*.*visualization*.*credible*.*compute_credible_interval*) [80].

We performed posterior predictive checks by simulating CNV dynamics using the MAP estimate as well as 50 parameter values sampled from the posterior distribution. We calculated root mean square error (RMSE) and correlation to measure agreement of the 50 posterior predictions with the observation, and report the mean and 95% confidence intervals for these measures. For inference on sets of observations, we calculated the RMSE and correlation coefficient between the posterior predictions and each of the three held out observations, and report the mean and 95% confidence intervals for these measures over all three held out observations.

We calculated *Akaike’s information criteria* (AIC) using the standard formula

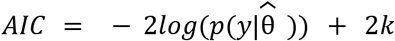

where 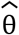 is the MAP estimate, *k* = 2 is the number of inferred parameters, *y* is the observed data, and *p* is the inferred posterior distribution. We calculated *Watanabe-Akaike information criterion* or *widely applicable information criterion* (WAIC) according to both commonly used formulas:

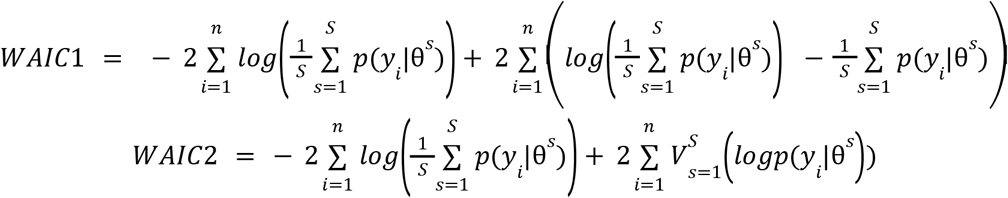

where *S* is the number of draws from the posterior distribution, θ^*s*^ is a sample from the posterior, and 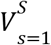 is the posterior sample variance.

### Pairwise competitions

We isolated CNV-containing clones from the populations on the basis of fluorescence, and performed pairwise competitions between each clone and an unlabeled ancestral (FY4) strain. We also performed competitions between the ancestral *GAP1* CNV reporter strain, with and without barcodes. To perform the competitions, we grew fluorescent *GAP1* CNV clones and ancestral clones in glutamine-limited chemostats until they reached steady state [61]. We then mixed the fluorescent strains with the unlabeled ancestor in a ratio of approximately 1:9, and performed competitions in the chemostats for 92 hours or about 16 generations, sampling approximately every 2-3 generations. For each time point, at least 100,000 cells were analyzed using an Accuri flow cytometer to determine the relative abundance of each genotype. Previously, we established that the ancestral *GAP1* CNV reporter has no detectable fitness effect compared to the unlabeled ancestral strain [61]. However, the *GAP1* CNV reporter with barcodes does appear to have a slight fitness cost associated with it, therefore, we took slightly different approaches to determine the selection coefficient relative to the ancestral state depending on whether or not a *GAP1* CNV containing clone was barcoded. If a clone was not barcoded, we detemined relative fitness using linear regression of the natural logarithm of the ratio of the two genotypes against the number of elapsed hours. If a clone was barcoded, relative fitness using linear regression of the natural logarithm of the ratio of the barcoded *GAP1* CNV containing clone to the unlabeled ancestor, and the natural logarithm of the ratio of the unevolved barcoded *GAP1* CNV reporter ancestor to the unlabeled ancestor against the number of elapsed hours, adding an additional interaction term for the evolved versus ancestral state. We converted relative fitness from per hour to generation by dividing by the natural log of two.

### Barcode sequencing

In our prior study, populations with lineage tracking barcodes and the *GAP1* CNV reporter were evolved in glutamine-limited chemostats [61], and whole population samples were periodically frozen in 15% glycerol. To extract DNA, we thawed pelleted cells using centrifugation and extracted genomic DNA using a modified Hoffman-Winston protocol, preceded by incubation with zymolyase at 37°C to enhance cell lysis [106]. We measured DNA quantity using a fluorometer, and used all DNA from each sample as input to a sequential PCR protocol to amplify DNA barcodes which were then purified using a Nucleospin PCR clean-up kit, as described previously[61,100].

We measured fragment size with an Agilent TapeStation 2200 and performed qPCR to determine the final library concentration. DNA libraries were sequenced using a paired-end 2 × 150 bp protocol on an Illumina NovaSeq 6000 using an XP workflow. Standard metrics were used to assess data quality (Q30 and %PF). We used the Bartender algorithm with UMI handling to account for PCR duplicates and to cluster sequences with merging decisions based solely on distance except in cases of low coverage (<500 reads/barcode), for which the default cluster merging threshold was used [69]. Clusters with a size less than 4 or with high entropy (>0.75 quality score) were discarded. We estimated the relative abundance of barcodes using the number of unique reads supporting a cluster compared to total library size. Raw sequencing data is available through the SRA, BioProject ID PRJNA767552.

### Detecting adaptive lineages in barcoded clonal populations

To detect spontaneous adaptive mutations in a barcoded clonal cell population that is evolved for over time, we used a Python-based pipeline (Li and Sherlock, in prep; https://github.com/FangfeiLi05/PyFitMut) based on a previously developed theoretical framework (Levy et al., 2015). The pipeline identifies adaptive lineages and infers their fitness effects and establishment time.. In a barcoded population, a lineage refers to cells that share the same DNA barcode. For each lineage in the barcoded population, beneficial mutations continually occur at a total beneficial mutation rate Ub, with fitness effect s, with a certain spectrum of fitness effects of mutations —(s). If a beneficial mutant survives random drift and becomes large enough to grow deterministically (exponentially), we say that the mutation carried by the mutant has established. Here, we use Wrightian fitness s, which is defined as average number of additional t offspring of a cell per generation, that is, n(t) = n(0)·(1 + s), with n(t) being the total number of cells at generation t (can be non-integers). Briefly, for each lineage, assuming that the lineage is adaptive (i.e., a lineage with a beneficial mutation occurred and established), then estimates of the fitness effect and establishment time of each lineage are made by random initialization, and the expected trajectory of each lineage is estimated and compared to the measured trajectory. Fitness effect and establishment time estimates are iteratively adjusted to better fit the observed data until an optimum is reached. At the same time, the expected trajectory of the lineage is also estimated assuming that the lineage is neutral. Finally, Bayesian inference is used to determine whether the lineage is adaptive or neutral. An accurate estimation of the mean fitness is necessary to detect mutations and quantify their fitness effects, but the mean fitness is a quantity that cannot be measured directly from the evolution. Rather, it needs to be inferred through other variables. Previously, the mean fitness was estimated by monitoring the decline of neutral lineages (Levy et al., 2015). However, this method fails when there is an insufficient number of neutral lineages as a result of low sequencing read depth. Here, we instead estimate the mean mean fitness using an iterative method. Specifically, we first initialize the mean fitness of the population as zero at each sequencing time point, then we estimate the fitness effect and establishment time for adaptive mutations, then we recalculate the mean fitness with the optimized fitness and establishment time estimates, repeating the process for several iterations until the mean fitness converges. We established the improved the accuracy of the method using simulated data (Li and Sherlock, in prep).

## Acknowledgements

We thank Uri Obolski, Ilia Kohanovski, Mark Siegal, and Molly Przeworski for discussions and comments. This work was supported in part by Israel Science Foundation (YR 552/19) and Minerva Stiftung Center for Lab Evolution (YR), grants from the NIH (R01 GM134066) and NSF (MCB1818234) (DG), grants from the NIH (R35 GM131824 and R01 AI136992) (GS), and an NSF GRFP (DGE1342536) (GA).

## Supplement

### Supplementary Files. Assessing inference method performance on single experiments

This is a zip folder containing the results of inference on single observations. Each file in the folder is named with the following naming convention: Model_Method_FlowType_SimulationBudget_InferenceSet_all.pdf. Each file contains 20 pages, each page corresponding to one of the 20 simulated synthetic observations. For NPE, each file corresponds to one training set (each observation was evaluated on a single amortized posterior was used) each page contains 6 panels, from top left to bottom right: a description of parameters used to generate the synthetic observation, a plot of the synthetic observation, the marginal posterior distribution for CNV formation rate, a plot of the posterior predictive check, the joint posterior distribution, and the marginal posterior distribution for CNV selection coefficient. For ABC-SMC, each page contains 8 panels from top left to bottom right: a description of parameters used to generate the synthetic observation, a plot of the synthetic observation, the effective sample size for each iteration of inference, epsilon values for each iteration of inference, the marginal posterior distribution for CNV formation rate for each iteration of inference, a plot of the posterior predictive check, the final joint posterior distribution, and the marginal posterior distribution for CNV selection coefficient for each iteration of inference. When the starting particle size = 100, the simulation budget was 10,000; when starting particle size = 1000, the simulation budget was 100,000.

**Supplementary Figure 1.**
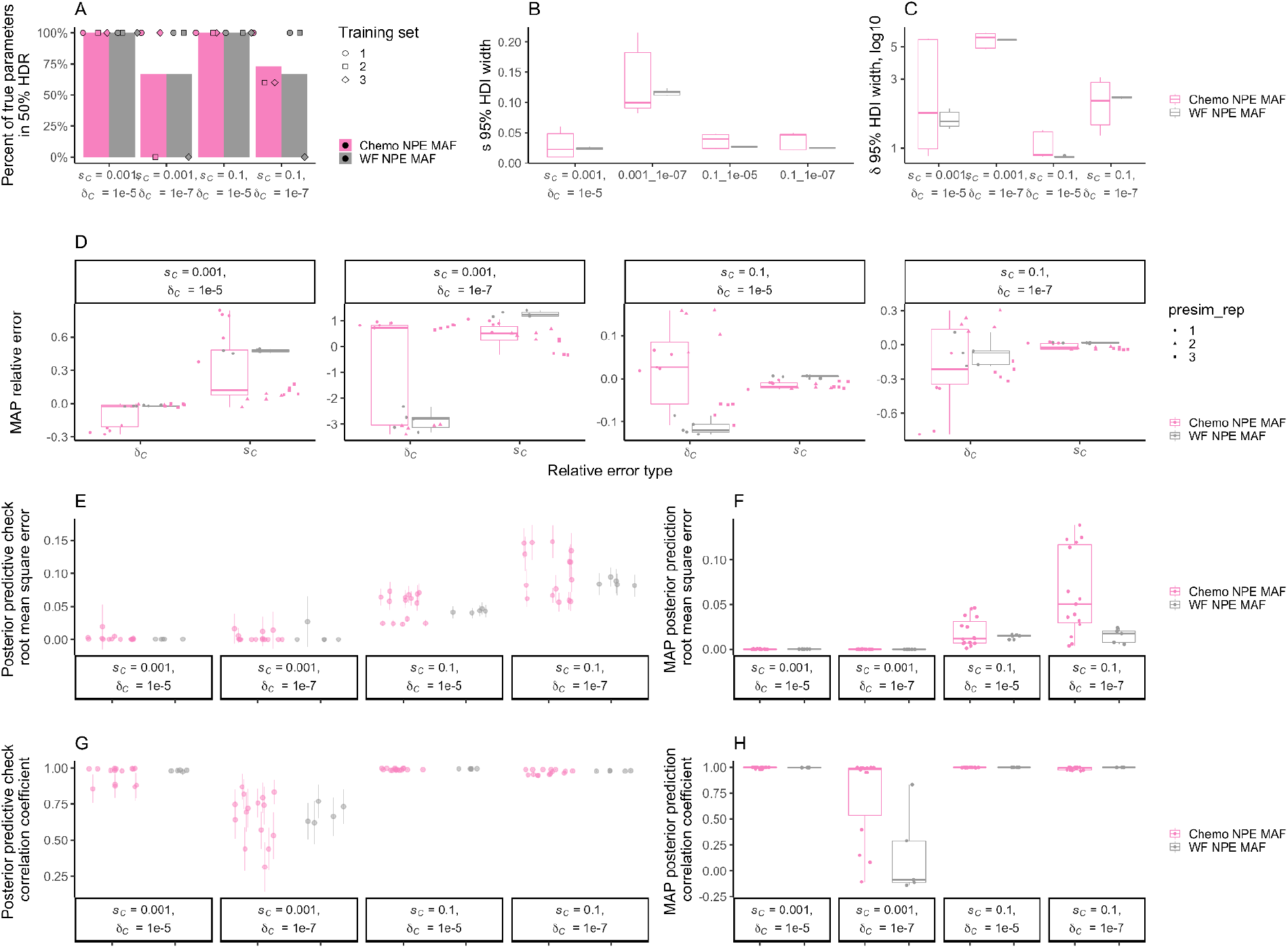
Performance assessment of NPE with MAF using single simulated synthetic observations. These show the results of inference on five simulated synthetic observations using either the Wright-Fisher (WF) or chemostat (Chemo) model per combination of fitness effect *s*_*C*_ and mutation rate *δ*_C_. Here we show the results of performing one training set with NPE with MAF using 100,000 simulations for training and using the same amortized network to infer a posterior for each replicate synthetic observation. **A)** Percentage of true parameters within the 50% HDR. **B)** Distribution of widths of the fitness effect *s*_*C*_ 95% highest density interval (HDI) calculated as the difference between the 97.5 percentile and 2.5 percentile, for each inferred posterior distribution. **C)** Distribution of the number of orders of magnitude encompassed by the mutation rate *δ* 95% HDI, calculated as difference of the base 10 logarithms of the 97.5 percentile and 2.5 percentile, for each inferred posterior distribution. **D)** Log ratio MAP estimate as compared to true parameters for *s*_*C*_, *δ*_C_, and total relative error. Note that each panel has a different y axis. **E)** Mean and 95% confidence interval for RMSE of 50 posterior predictions as compared to the synthetic observation for which inference was performed. **F)** RMSE of posterior prediction generated with MAP parameters as compared to the synthetic observation for which inference was performed. **G)** Mean and 95% confidence interval for correlation coefficient of 50 posterior predictions compared to the synthetic observation for which inference was performed. **H)** Correlation coefficient of posterior prediction posterior prediction generated with MAP parameters compared to the synthetic observation for which inference was performed.

**Supplementary Figure 2.**
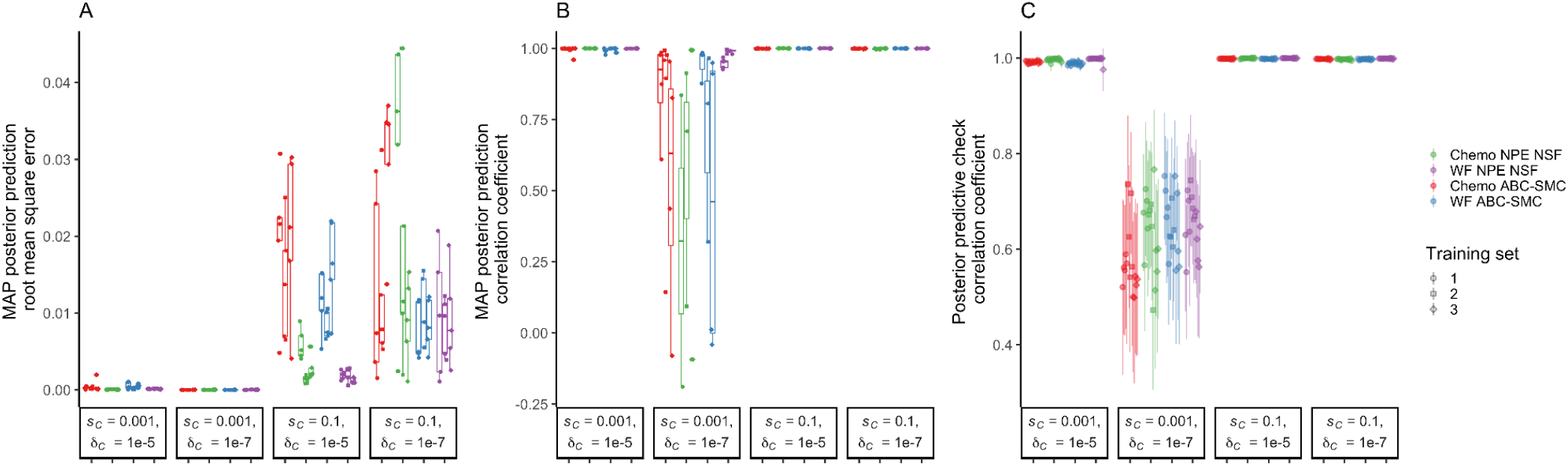
NPE with the Wright-Fisher model performs as well or better than other combinations of model and method. Results of inference on five simulated single synthetic observations using either the Wright-Fisher (WF) or chemostat (Chemo) model per combination of fitness effect *s*_*C*_ and mutation rate *δ*_C_. Here we show the results of performing training with NPE with NSF using 100,000 simulations for training and using the same amortized network to infer a posterior for each replicate synthetic observation, or ABC-SMC when the training budget was 100,000. **A)** RMSE (lower is better) of posterior prediction generated with MAP parameters as compared to the synthetic observation on which inference was performed. **B)** Correlation coefficient (higher is better) of posterior prediction generated with MAP parameters compared to the synthetic observation on which inference was performed. **C)** Mean and 95% confidence interval for correlation coefficient (higher is better) of 50 posterior predictions (sampled from the posterior distribution) compared to the synthetic observation on which inference was performed.

**Supplementary Figure 3.**
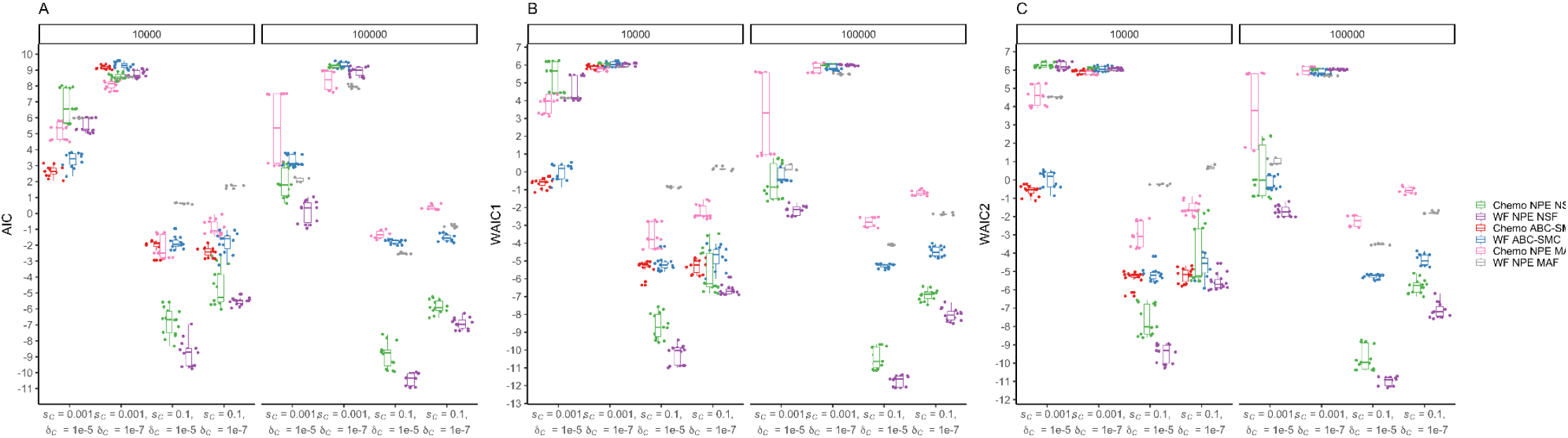
NPE and WF have the lowest information criteria. WAIC and AIC (lower is better) of models fitted on single synthetic observations using either the Wright-Fisher (WF) or chemostat (Chemo) model and either ABC-SMC or NPE for different combinations of fitness effect *s*_*C*_ and mutation rate *δ*_C_ with simulation budgets of 10,000 or 100,000 simulations per inference procedure (facets). We were unable to complete ABC-SMC with the chemostat model (red) when the training budget was 100,000 within a reasonable time frame.

**Supplementary Figure 4.**
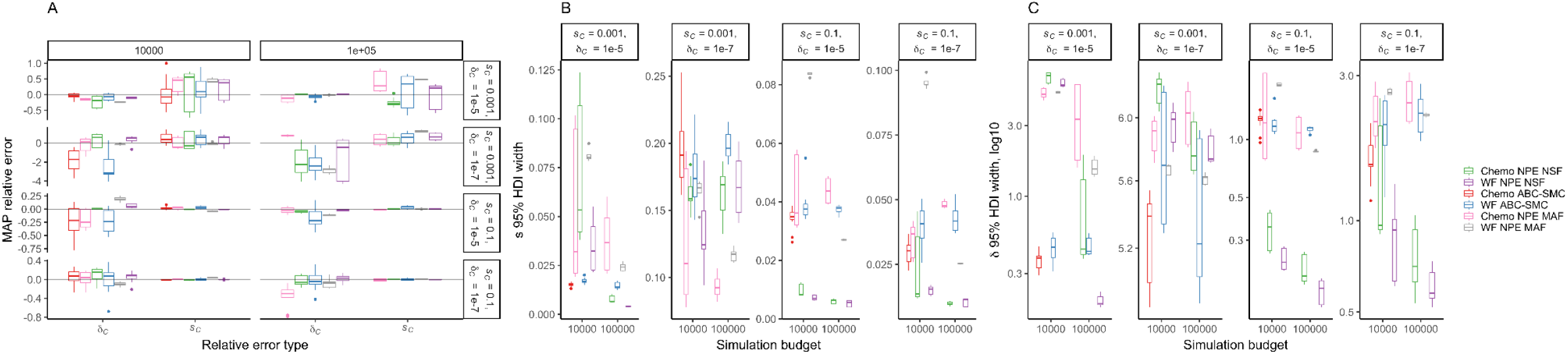
Effect of simulation budget on relative error of MAP estimate and width of HDIs for inference on single synthetic observations. The grey line in (**A**) indicates a relative error of zero (i.e., no difference between MAP parameters and true parameters). We were unable to complete ABC-SMC with the chemostat model (red) when the training budget was 100,000 within a reasonable time frame.

**Supplementary Figure 5.**
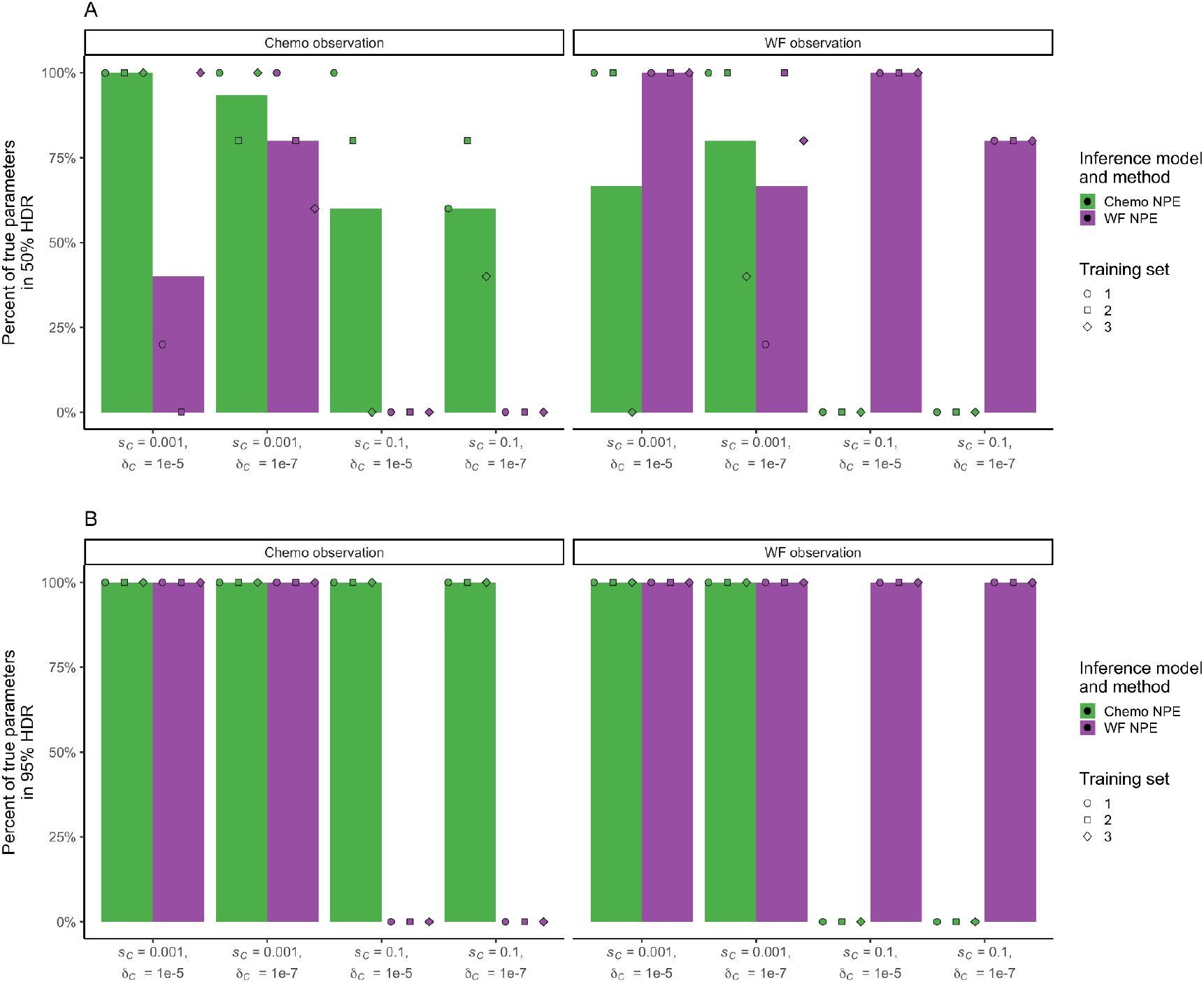
Results of inference on five simulated synthetic observations generated using either the Wright-Fisher (WF) or chemostat (Chemo) model per combination of fitness effect *s*_*C*_ and mutation rate *δ*_C_. We performed inference on each synthetic observation using both models. For NPE, each training set corresponds to an independent amortized posterior trained with 100,000 simulations, with which each synthetic observation was evaluated. **A**) Percentage of true parameters within the 50% HDR. The bar height shows the average of three training sets. **B**) Percentage of true parameters within the 95% HDR. The bar height shows the average of three training sets.

**Supplementary Figure 6.**
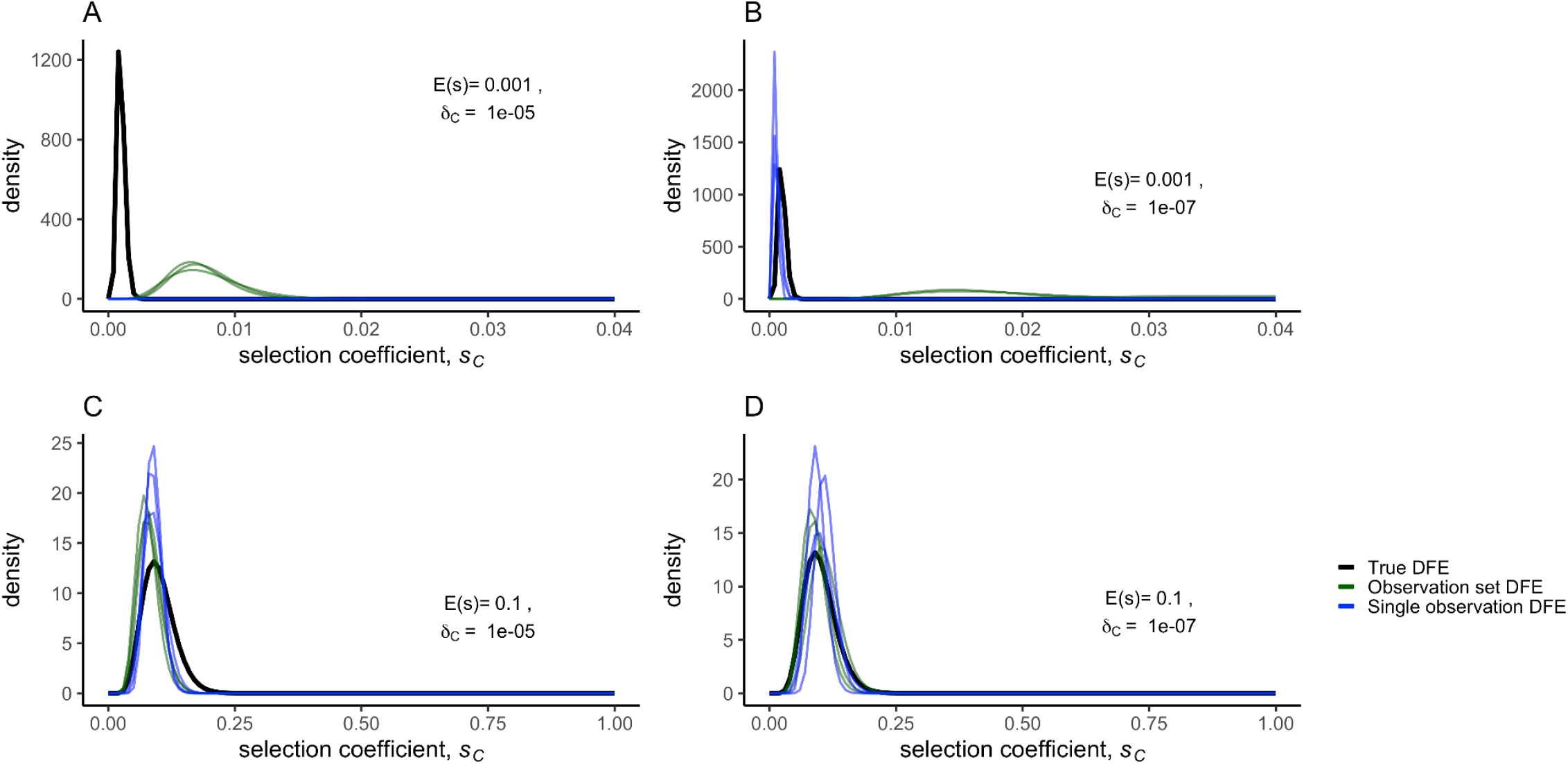
A set of eleven simulated synthetic observations was generated from a Wright-Fisher model with CNV selection coefficients sampled from an Gamma distribution where *α* = 10 of fitness effects (DFE) (black curve). The MAP DFEs (blue curves) were directly inferred using three different subsets of eight out of eleven synthetic observations. We also inferred the selection coefficient for each observation in the set of eleven individually, and fit Gamma distributions to sets of eight inferred selection coefficients (green curves). All inferences were performed with NPE using the same amortized network to infer a posterior for each set of eight synthetic observations or each single observation.

**Supplementary Table 1.** Kullback–Leibler divergence for Gamma distributions fit from single inferred selection coefficients versus the true underlying DFE, or for directly inferred Gamma distributions versus the true underlying DFE.

| KL divergence:<br>Gamma fit from<br>single inferred $s_c$ | KL divergence: $\alpha$<br>and $\beta$ directly<br>inferred from set of<br>observations | Observation set name | True $\alpha$ | True $\beta$ | True $\delta_C$ |
| --- | --- | --- | --- | --- | --- |
| 1359.8 | 572810.04 | WF_shape1_scale0.001_mut5 | 1.0 | 0.001 | 1E-05 |
| 856459.8 | 200872.69 | WF_shape1_scale0.001_mut5 | 1.0 | 0.001 | 1E-05 |
| 1338.0 | 644533.9 | WF_shape1_scale0.001_mut5 | 1.0 | 0.001 | 1E-05 |
| 38967.7 | 664134.98 | WF_shape1_scale0.001_mut7 | 1.0 | 0.001 | 1E-07 |
| 6522383.1 | 854560.75 | WF_shape1_scale0.001_mut7 | 1.0 | 0.001 | 1E-07 |
| 38652.8 | 372597.74 | WF_shape1_scale0.001_mut7 | 1.0 | 0.001 | 1E-07 |
| 233.8 | 1200.59 | WF_shape1_scale0.1_mut5 | 1.0 | 0.1 | 1E-05 |
| 230.7 | 220.63 | WF_shape1_scale0.1_mut5 | 1.0 | 0.1 | 1E-05 |
| 233.0 | 1161.7 | WF_shape1_scale0.1_mut5 | 1.0 | 0.1 | 1E-05 |
| 9.4 | 5151.41 | WF_shape1_scale0.1_mut7 | 1.0 | 0.1 | 1E-07 |
| 6.6 | 1255.33 | WF_shape1_scale0.1_mut7 | 1.0 | 0.1 | 1E-07 |
| 21.7 | 1130.75 | WF_shape1_scale0.1_mut7 | 1.0 | 0.1 | 1E-07 |
| 2079309.0 | 381627.51 | WF_shape10_scale0.0001_mut5 | 10.0 | 0.0001 | 1E-05 |
| 1719636.6 | 562084.78 | WF_shape10_scale0.0001_mut5 | 10.0 | 0.0001 | 1E-05 |
| 2125542.6 | 543314.78 | WF_shape10_scale0.0001_mut5 | 10.0 | 0.0001 | 1E-05 |
| 32299.9 | 1124713.46 | WF_shape10_scale0.0001_mut7 | 10.0 | 0.0001 | 1E-07 |
| 133767.3 | 818178.69 | WF_shape10_scale0.0001_mut7 | 10.0 | 0.0001 | 1E-07 |
| 51454.3 | 993824.56 | WF_shape10_scale0.0001_mut7 | 10.0 | 0.0001 | 1E-07 |
| 336.3 | 123.01 | WF_shape10_scale0.01_mut5 | 10.0 | 0.01 | 1E-05 |
| 231.7 | 274.56 | WF_shape10_scale0.01_mut5 | 10.0 | 0.01 | 1E-05 |
| 74.1 | 134.88 | WF_shape10_scale0.01_mut5 | 10.0 | 0.01 | 1E-05 |
| 334.6 | 49.24 | WF_shape10_scale0.01_mut7 | 10.0 | 0.01 | 1E-07 |
| 228.3 | 25.18 | WF_shape10_scale0.01_mut7 | 10.0 | 0.01 | 1E-07 |
| 22.0 | 66.22 | WF_shape10_scale0.01_mut7 | 10.0 | 0.01 | 1E-07 |

**Supplementary Figure 7.**
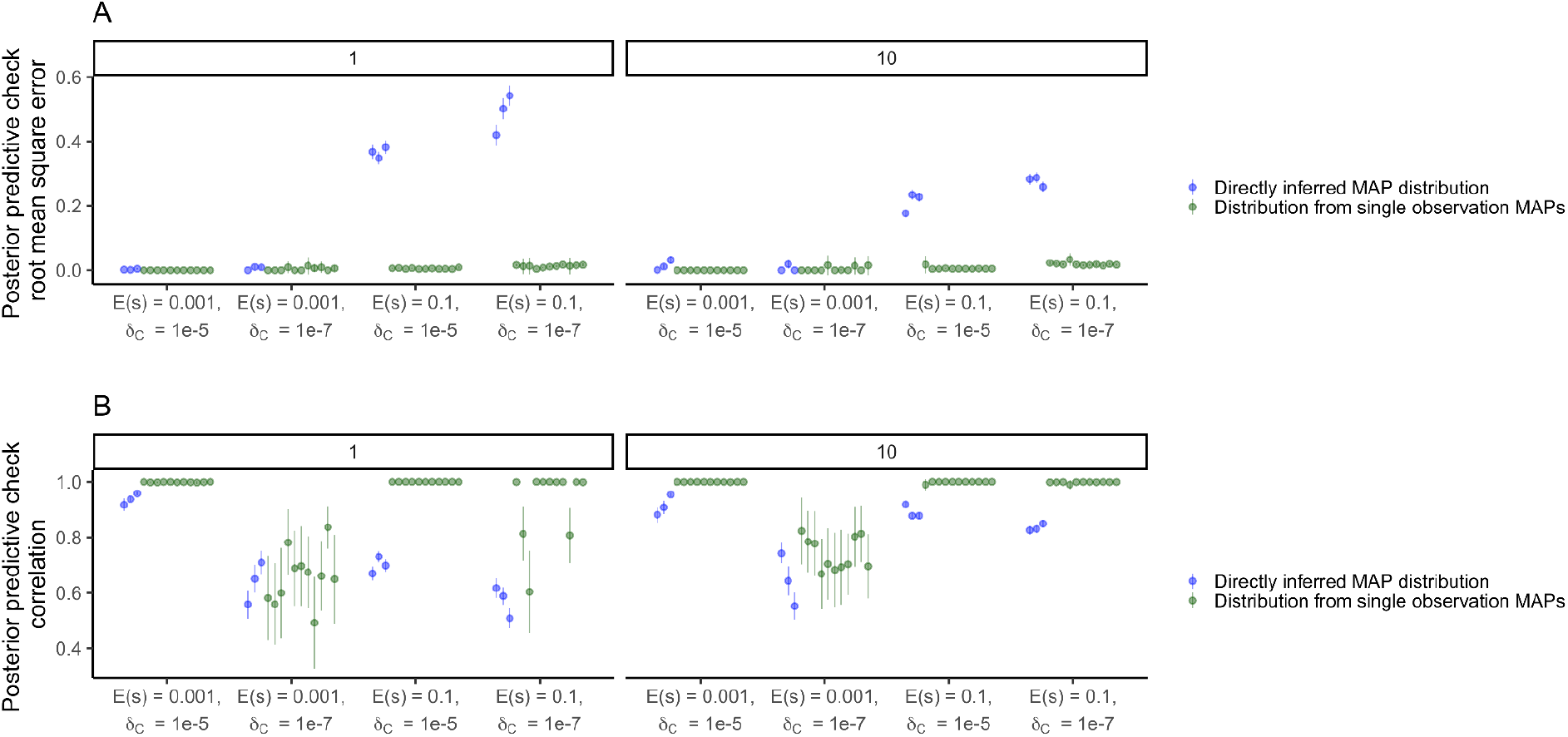
Out-of-sample posterior predictive accuracy using root mean square error (**A**) or correlation (**B**) using three held out observations when *α* and *β* are directly inferred from the other eight observations, for *α* = 1 or *α* = 10 (facets).

**Supplementary Figure 8.**
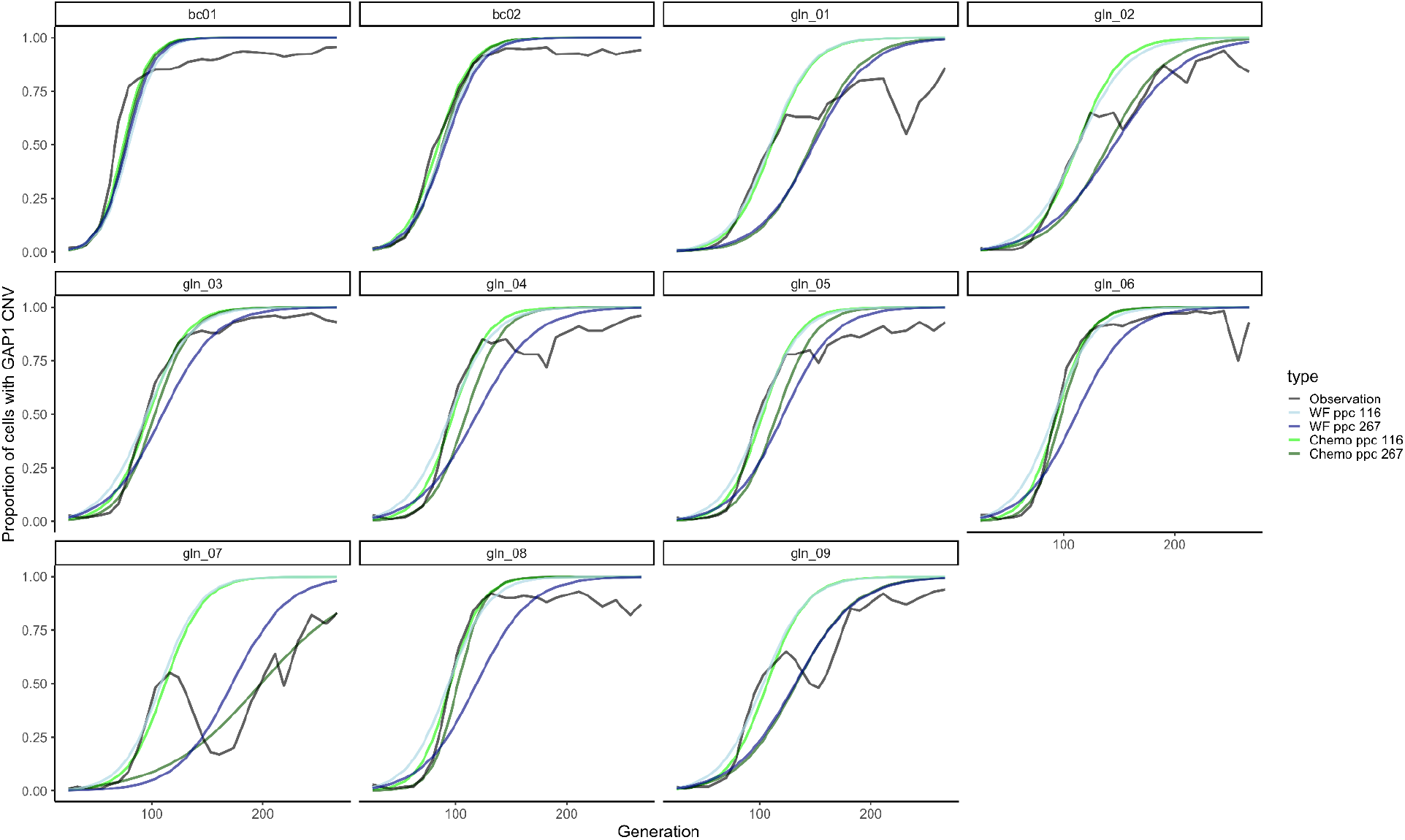
Proportion of the population with a *GAP1* CNV in the experimental observations (black) and in posterior predictions using the MAP estimate shown in panels A and B with either the Wright-Fisher (WF) or chemostat (Chemo) model. Inference was performed with all data up to generation 267 (WF ppc 267, Chemo ppc 267), or excluding data after generation 116 (WF ppc 116, Chemo ppc 116). Mutation rate and fitness effect of other beneficial mutations set to 10^−5^ and 10^−3^, respectively.

**Supplementary Figure 9.**
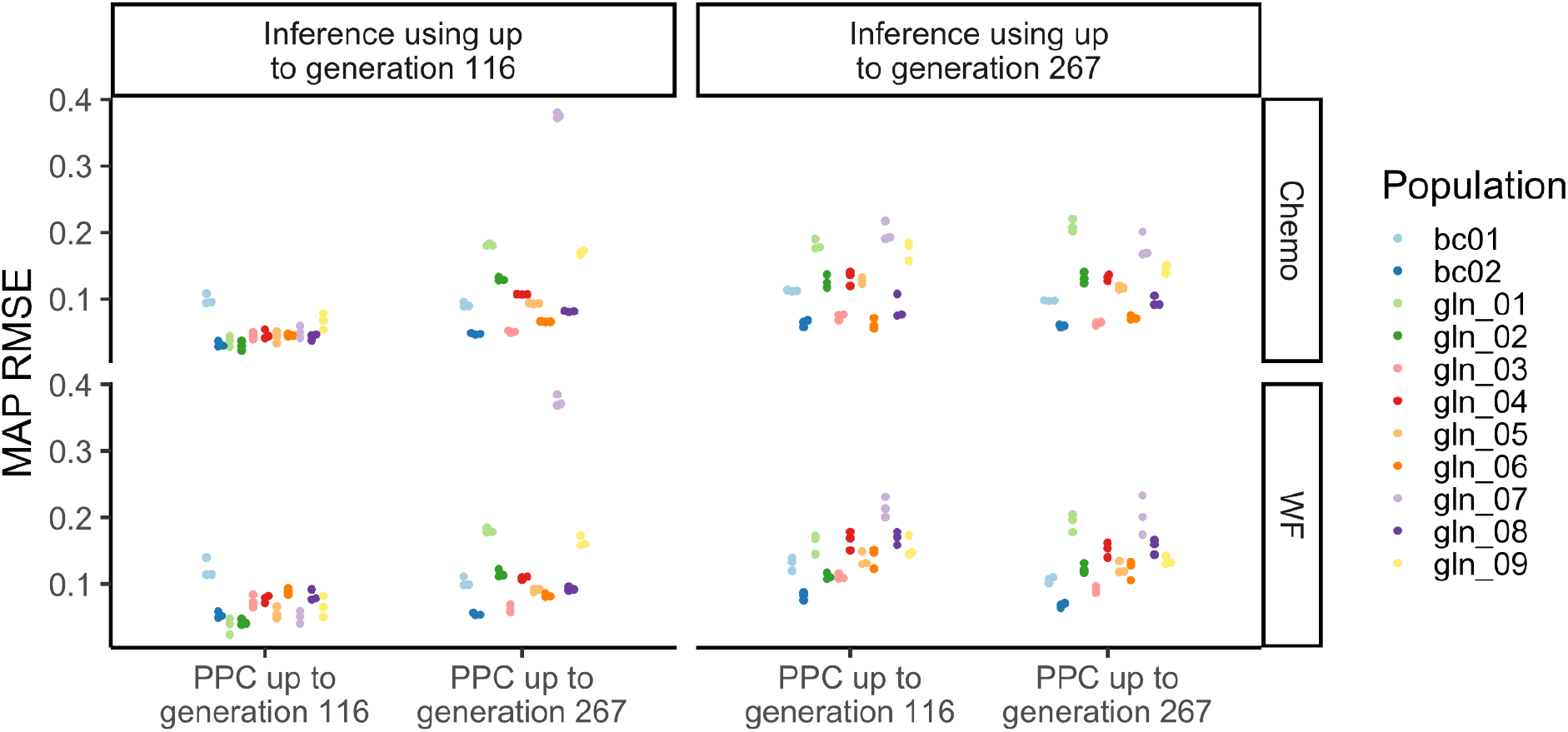
MAP posterior prediction root mean square error (RMSE) when inference was performed excluding data after generation 116 (left) or using all data up to generation 267 (right). RMSE was calculated using either the first 116 generations, or using up to generation 267 (x-axis).

**Supplementary Figure 10.**
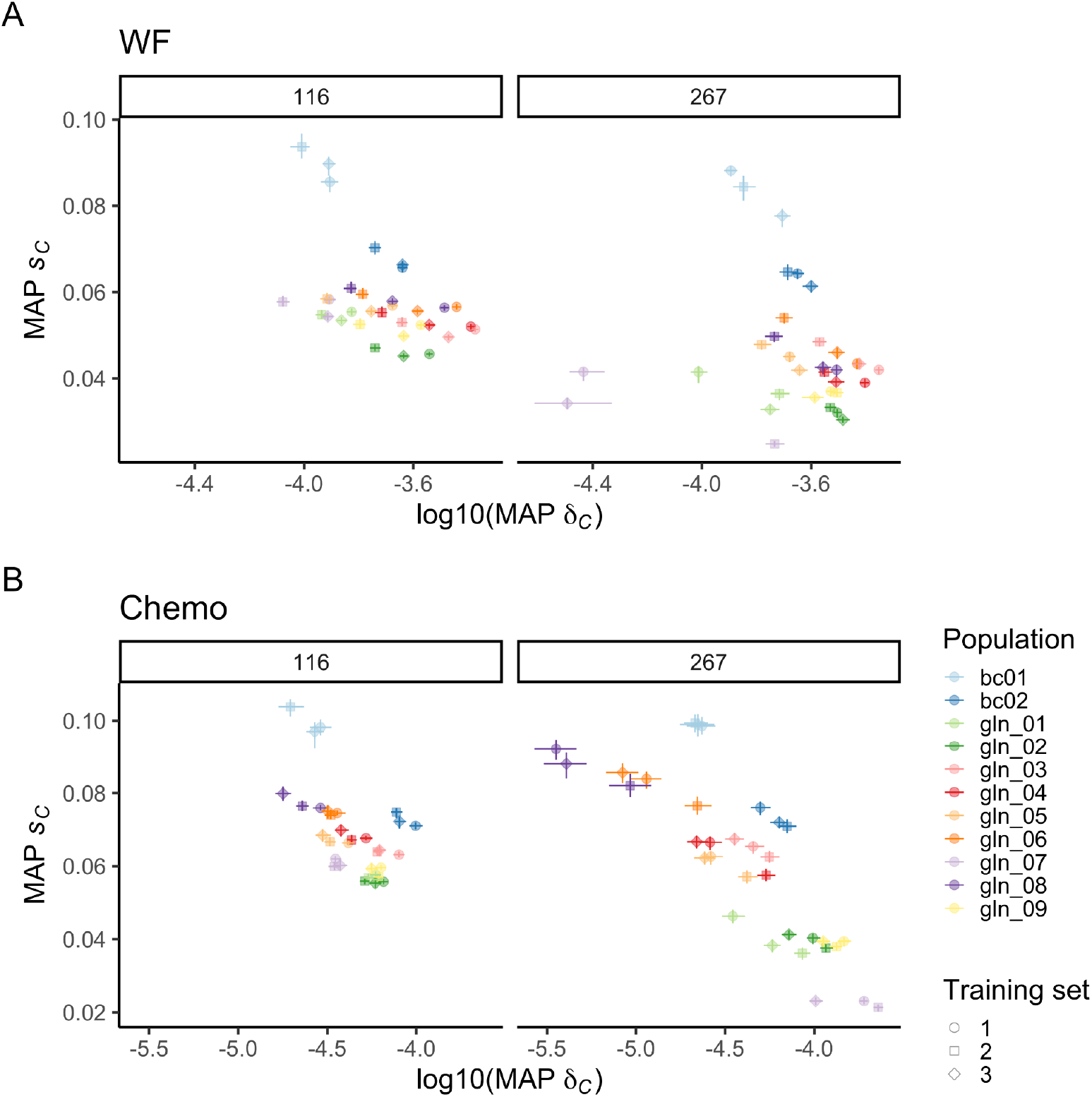
The inferred MAP estimate and 95% highest density intervals (HDI) for fitness effect *s*_*C*_ and formation rate *δ*_C_, using the **(A)** Wright-Fisher (WF) or **(B)** chemostat (Chemo) model and NPE for each experimental population from Lauer et al. (2018). Inference was either performed with data up to generation 116 or with all data, up to generation 267 (facets). Each training set corresponds to three independent amortized posterior distributions estimated with 100,000 simulations.

**Supplementary Figure 11.**
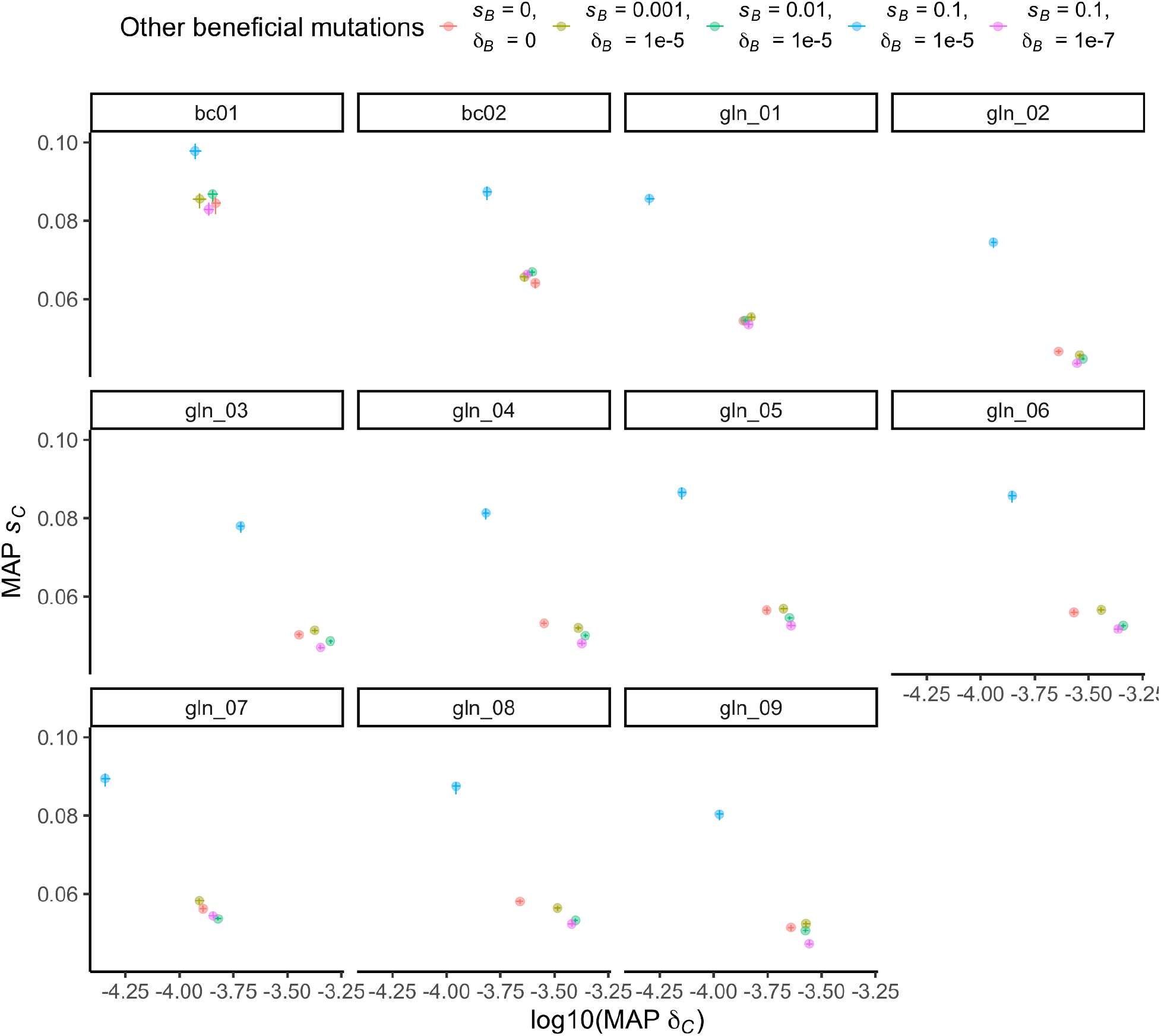
*GAP1* CNV formation rate and selection coefficient inferred using NPE with the Wright-Fisher model does not change dramatically when other beneficial mutations have different selection coefficients *s*_*B*_ and formation rates *δ*_B_, except when both *s*_*B*_ and *δ*_B_ are high (blue).

**Supplementary Figure 12.**
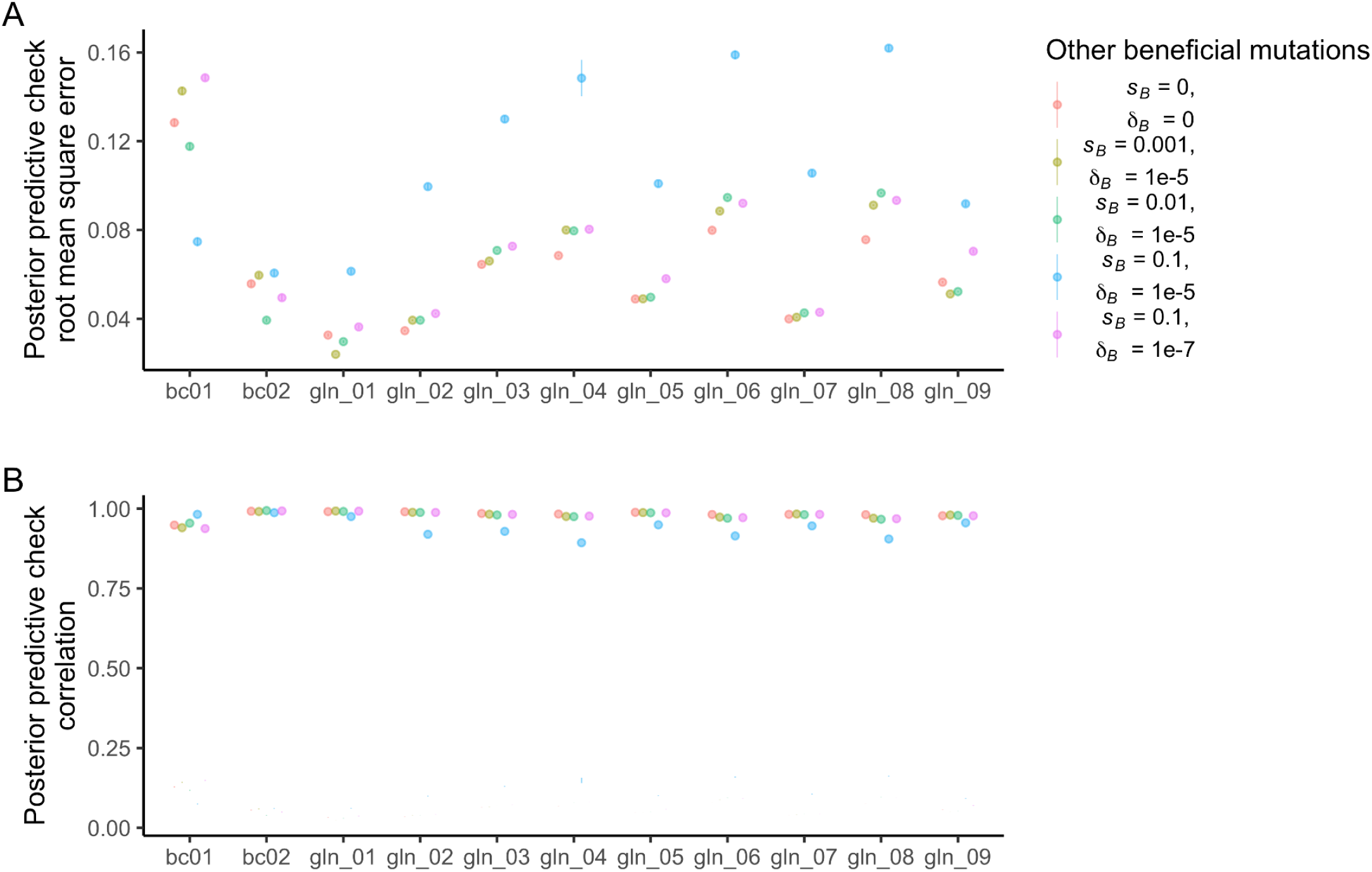
Mean and 95% confidence interval for RMSE (**A**) and correlation (**B**) of 50 posterior predictions compared to empirical observations up to generation 116, using posterior distributions inferred when other beneficial mutations have different selection coefficients *s*_*B*_ and formation rates *δ*_B_.

**Supplementary Figure 13.**
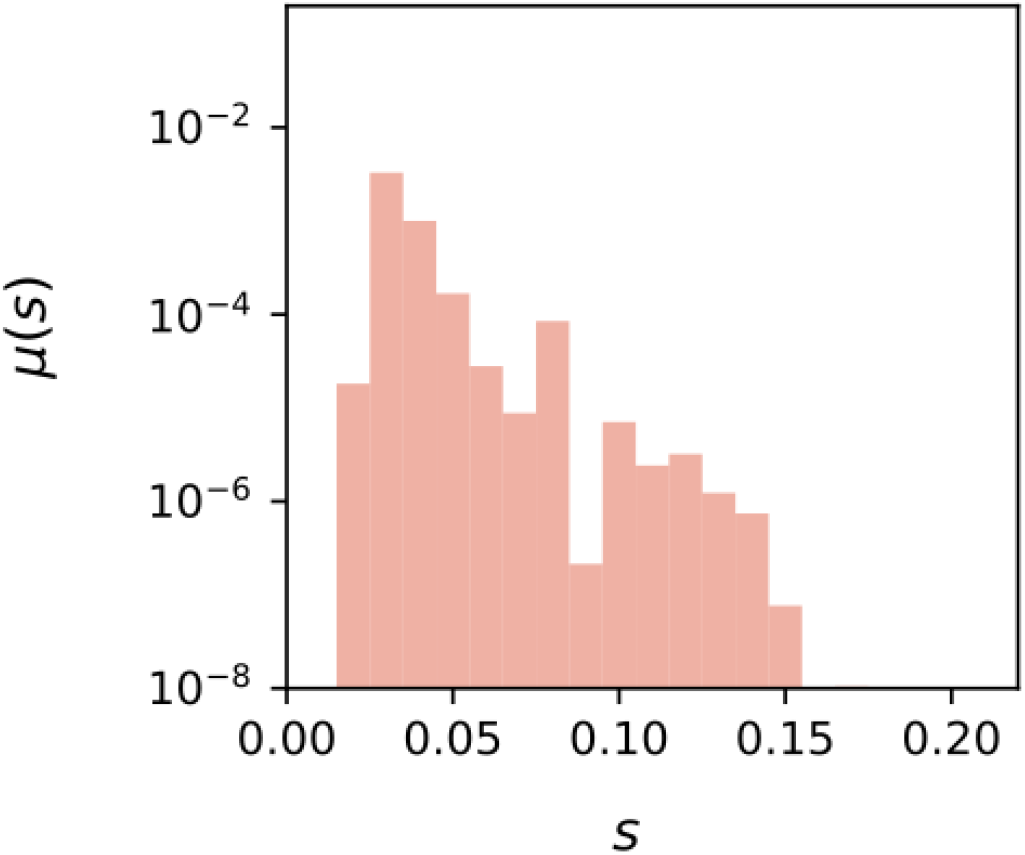
The spectrum of mutation rates, *μ(s)*, as a function of fitness effect, *s*, for all beneficial mutations, inferred from barcode sequencing of population bc01.

